# Histone modifications directly modulate transcription factor binding to nucleosomes

**DOI:** 10.64898/2026.10.21.760833

**Authors:** Luca Mariani, Martha L. Bulyk

**Author notes:** Corresponding authors. (L.M.); (M.L.B.).

## Abstract

Transcription factors (TFs) that can recognize specific DNA sequences within nucleosomes play key roles in regulating cell fate transitions, but what regulates their condition-dependent binding to nucleosomes remains poorly understood. Here, we determined the direct effects of H3K27ac, H3K27me3 and H3K9me3 on binding by 7 human TFs to a highly diverse library of nucleosomes. We found that these marks influence the preference of FOXA1 and PAX7 to bind as monomers versus dimers to nucleosomes. Furthermore, H3K27ac impairs TF binding at internal regions of nucleosomes, enhancing end-binding preferences, for most of the assayed TFs. Our results suggest that histone marks represent an additional layer of *cis*-regulatory encoding that directly regulates TF interactions with nucleosomal binding sites.

---

In eukaryotes, gene expression programs are regulated dynamically by the binding of transcription factors (TFs) to sequence-specific DNA binding sites in chromatin, but whether and how TFs recognize their sites in regions occluded by nucleosomes remains unclear. A heterogenous group of “pioneer” TFs can bind to inaccessible *cis*-regulatory elements for subsequent transcriptional activity, therefore serving as gatekeepers to cellular differentiation (*1, 2*). Pioneers bind nucleosomal sites in a cell-type or developmental stage-specific manner (*e.g.*, Zelda (*3*), FOXA2, GATA4, OCT4 (*4, 5*)). Results from several *in vivo* studies indicate that pioneer binding to genomic target sites is associated with post-translational modifications (PTMs) on the unstructured tails of histones (*6*). Computational analyses of ENCODE ChIP-Seq data have identified associations of TF genomic occupancy with histone PTMs (*7, 8*) but have not elucidated the causal relationships nor the mechanisms underlying those associations.

Notably, biochemical studies of TF binding to nucleosomes typically are performed using unmodified histones (*9-13*). The histones’ lysine-rich tails can establish fuzzy interactions with the negatively charged DNA, potentially affecting nucleosome structure and recruitment of TFs (*14*). Indeed, histone tails have been shown *in vitro* to inhibit TF binding to nucleosomes, especially to the more internal regions of nucleosomes (*15*). A recent study reporting a cryo-EM structure of nucleosomes containing a synthetic DNA sequence bound by the DNA binding domain (DBD) of SOX2 showed that when bound to specific locations within the nucleosome, the SOX DBD repositions the N-terminal tail of histone 4 (H4) (*16*), indicating that they can interact in the absence of chromatin modifiers.

Both *in vitro* experiments and *in silico* molecular simulations have indicated that histone PTMs can decrease interactions between histone tails and DNA. H3 acetylation, which marks active chromatin, neutralizes the positive charge of the lysines on the H3 tail, impairing its interaction with DNA, thus causing an increase in nucleosome breathing (*17-20*) and α-helical content (*21*). Trimethylation of H3K9 and H3K27, which mark repressed chromatin, create hydrophobic pockets around the lysines, thus favoring H3 tail release from the nucleosome (*22*). A recent study that found direct interaction of H3K27 with the OCT4 DBD bound to linker DNA also revealed that both H3K27ac and H3K27me3 enhanced OCT4 cooperative binding to *LIN28B* and *nMATN1* nucleosomes, mostly favoring internal regions of the nucleosomes away from the dyad (*23*). However, whether a broader set of histone PTMs directly modulates interactions between additional TFs and their nucleosomal DNA binding sites remains unknown.

We previously developed PIONEAR-seq (*13*), a high-throughput biochemical assay to evaluate the binding specificity and positional preferences of TFs to nucleosome core particles (NCPs) assembled on known nucleosome positioning sequences (*i.e.*, *NRCAM*, *CX3CR1*, *ALB1*, Widom 601 (W601)) (*24*), across which we tiled short (20-bp) fully degenerate sequences representing a highly diverse repertoire of potential TF binding sites. PIONEAR-seq data revealed that, while TFs exhibit the same DNA sequence specificity in binding to free and nucleosomal DNA, TFs’ positional preferences of binding within nucleosomes are regulated by local contributions of DNA bendability (*13*). Here, we leveraged PIONEAR-seq to investigate the direct effects of histone PTMs on TF binding to nucleosomes assembled with H3 histones engineered to contain either H3K27ac (*25*) or analogs of H3K27me3 or H3K9me3 (*26*); methyl-lysine analogs within histones have served as a powerful tool (*27*) in numerous prior studies of modified nucleosomes, including their interactions with chromatin factors (*e.g.*, (*28, 29*)). Our analysis of the binding of 7 human pioneer TFs to these modified nucleosomes revealed PTM-induced modulation of TF binding, suggesting an additional *cis*-regulatory layer controlling TF binding to genomic recognition sites.

## TFs bind with comparable levels of specificity to modified as unmodified nucleosomes

To assess whether TFs that bind recognition sequences within nucleosomal DNA directly sense the presence of histone PTMs, we adapted PIONEAR-seq technology (*13*) by incorporating H3 histones containing PTMs that mark regulatory chromatin states (*30*), namely H3K27ac for active chromatin, H3K27me3 for facultative heterochromatin and H3K9me3 for constitutive heterochromatin (Fig. 1A). Briefly, to investigate the positional preferences of TF binding within nucleosomes, we tiled three input 147-bp nucleosomal positioning sequences (“templates”) – two genomic sequences (*CX3CR1, NRCAM*) and the high affinity, synthetic W601 sequence – with 20-bp random sequences (N_20_) and pooled them together. We reconstituted separate histone octamers containing each of the three different PTMs (*25, 26*) (fig. S1A) and mixed them with the highly diverse nucleosomal positioning sequence DNA libraries to assemble NCPs (fig. S1B). We then incubated each of the four different unmodified or modified NCP libraries separately with each of 7 human, full-length, GST-tagged TFs CEBPB, FOXA1, GATA1, GATA4, OCT4, PAX7, RUNX2) (*13*) that we expressed cell-free by *in vitro* transcription and translation (IVTT) and performed four consecutive rounds of selection for TF-bound nucleosomes (fig. S1C) (Methods). The DNA sequences from Round 3 and 4 libraries were separated by PCR according to their template of origin, thus creating 3 template-specific libraries (‘template replicates’) that were assayed in parallel in the same pool, then sequenced and analyzed for TF binding, as previously described (*13*). As negative controls, for each of the 7 TFs we performed analogous *in vitro* selections for binding to free DNA.

**Fig. 1.**
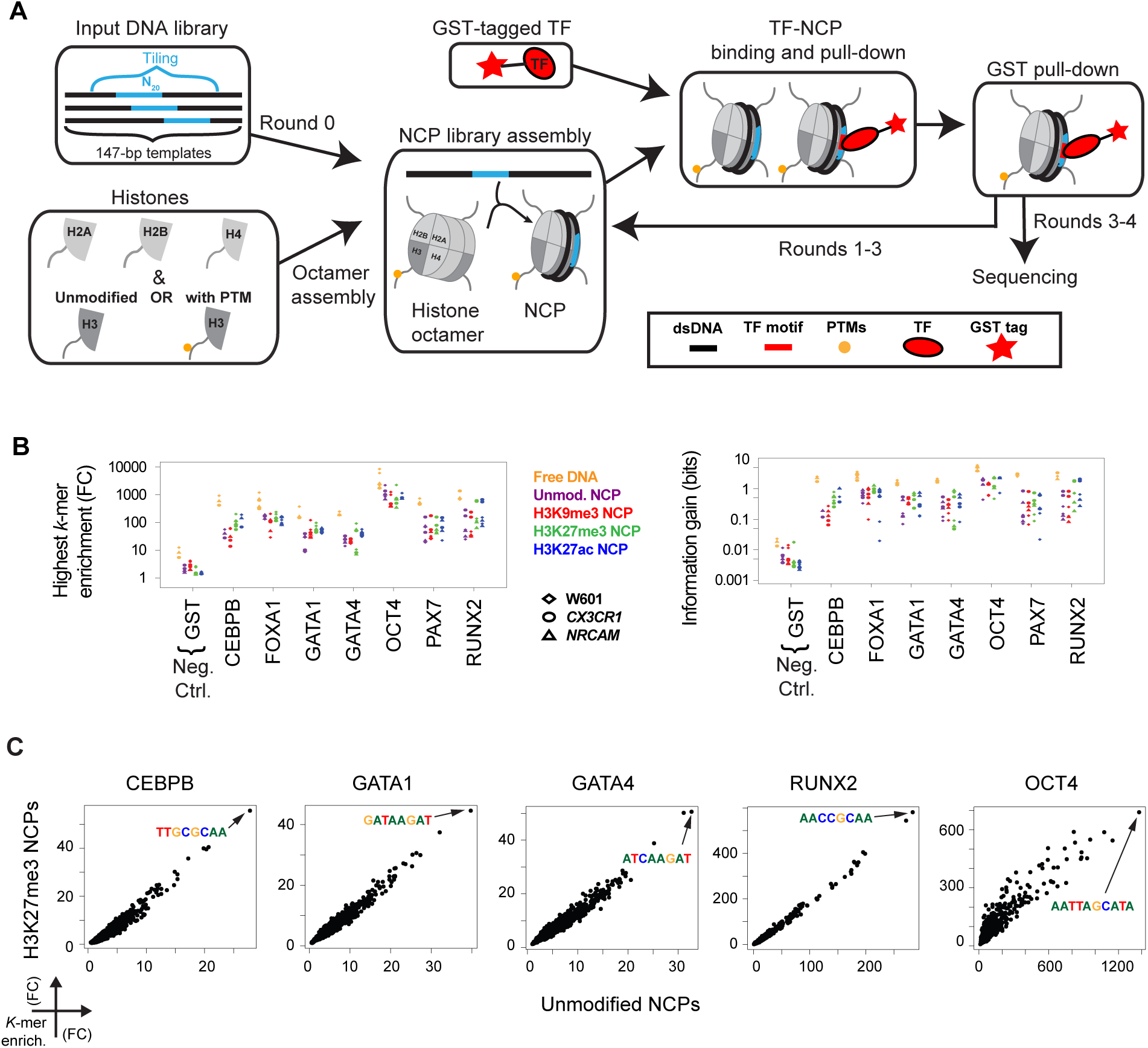
Incorporation of epigenetic histone modifications in PIONEAR-seq assays. **(A)** Workflow for integrating epigenetic histone marks into PIONEAR-seq assays (*13*) (Methods). **(B)** Highest *k*-mer enrichment *(left)* and information gain *(right)* in the PIONEAR-seq libraries stratified by template (symbols) and NCP type (colors). The enrichment of each *k*-mer is measured as the fold-change (FC) of its frequency in the PIONEAR-seq library versus the input library (see Methods). **(C)** Scatterplots of *k*-mer enrichment in *CX3CR1*-based NCPs assembled with either unmodified H3 (*x*-axis) or H3 trimethylated at lysine 27 (*y*-axis) and selected by the indicated TFs. For each TF, the most enriched *k*-mer is highlighted. The same patterns appear in libraries from the same TF-selected pool that were based on different templates (‘template replicates’) (fig. S2A, B, C) and for other NCP types (H3K9me3, H3K27ac) (fig. S3).

We quantified motif enrichment in each library using a *de novo k*-mer discovery method (*31*), from which we assigned to each TF an optimal *k*-mer binding site length, namely 8-mers for CEBPB, GATA1, GATA2, PAX7 and RUNX2, 9-mers for FOXA1, and 10-mers for OCT4, as in prior PIONEAR-seq analysis of these TFs (*13*). Motif selection in each library, as scored by the top *k*-mer enrichment (Fig. 1B, left panel) and by the entire library’s information gain (*31*) (Fig. 1B, right panel), showed that TFs exhibit comparable DNA binding specificity for the unmodified and modified nucleosomes. This specificity was significantly higher than that for a ‘GST alone’ negative control and less than for binding to free DNA, supporting prior studies that found TFs to recognize more degenerate (*32*) or shorter (*33*) motifs when bound to nucleosomes than to free DNA.

## Histone modifications modulate the sequence specificity of nucleosome binding by FOXA1 and PAX7

Comparison of the enriched *k*-mers from binding to modified versus unmodified NCPs showed very similar patterns of relative *k*-mer enrichment for 5 of the 7 TFs (CEBPB, GATA1, GATA4, OCT4, RUNX2) (Fig. 1C). The relative *k*-mer enrichment was similar across nucleosome sequence templates, modified and unmodified histones, and independent replicate PIONEAR-Seq assays (fig. S2, S3).

In contrast to CEBPB, GATA1, GATA4, OCT4 and RUNX2, however, the modified nucleosomes affected the DNA sequence preferences of FOXA1 and PAX7 (Fig. 2). FOXA1, which is a member of the FOX family characterized by a single winged helix-turn-helix forkhead (FKH) DBD, is a well-characterized pioneer factor in liver differentiation (*34*). As a monomer, FOX TFs recognize primarily the FKH sequence GTAAACA as well as a variety of alternative sequences (*35*). A study of FOXA1 genomic occupancy during liver differentiation (*32*) found that FOXA1 favors the FKH sequence on accessible chromatin, but more degenerate sequences, such as GTCAACA (DEG1) and GTMAATA (DEG2), on nucleosomes (Fig. 2A, left). In addition to FOXA1 monomeric binding, several motifs have been proposed for DNA-mediated FOXA1 homodimerization (*36*). Among them, the DIV0 motif, which comprises two divergent, overlapping DEG2 motifs (Fig. 2A, right), is known to be bound by FOXA1 in biochemical assays (*in vitro*) and to be involved in FOXA1 nucleosome binding in cellular assays (*in vivo*) near genes implicated in cancer progression (*37*). PIONEAR-Seq data revealed that the presence of PTMs increases FOXA1 recognition of DIV0, DEG2 and DEG1 sites, in contrast to strong preference for FKH sites on unmodified NCPs (Fig. 2B, C; fig. S4), suggesting that *in vivo* FOXA1 binds DIV0, DEG2 and DEG1 motifs preferentially on modified nucleosomes.

**Fig. 2.**
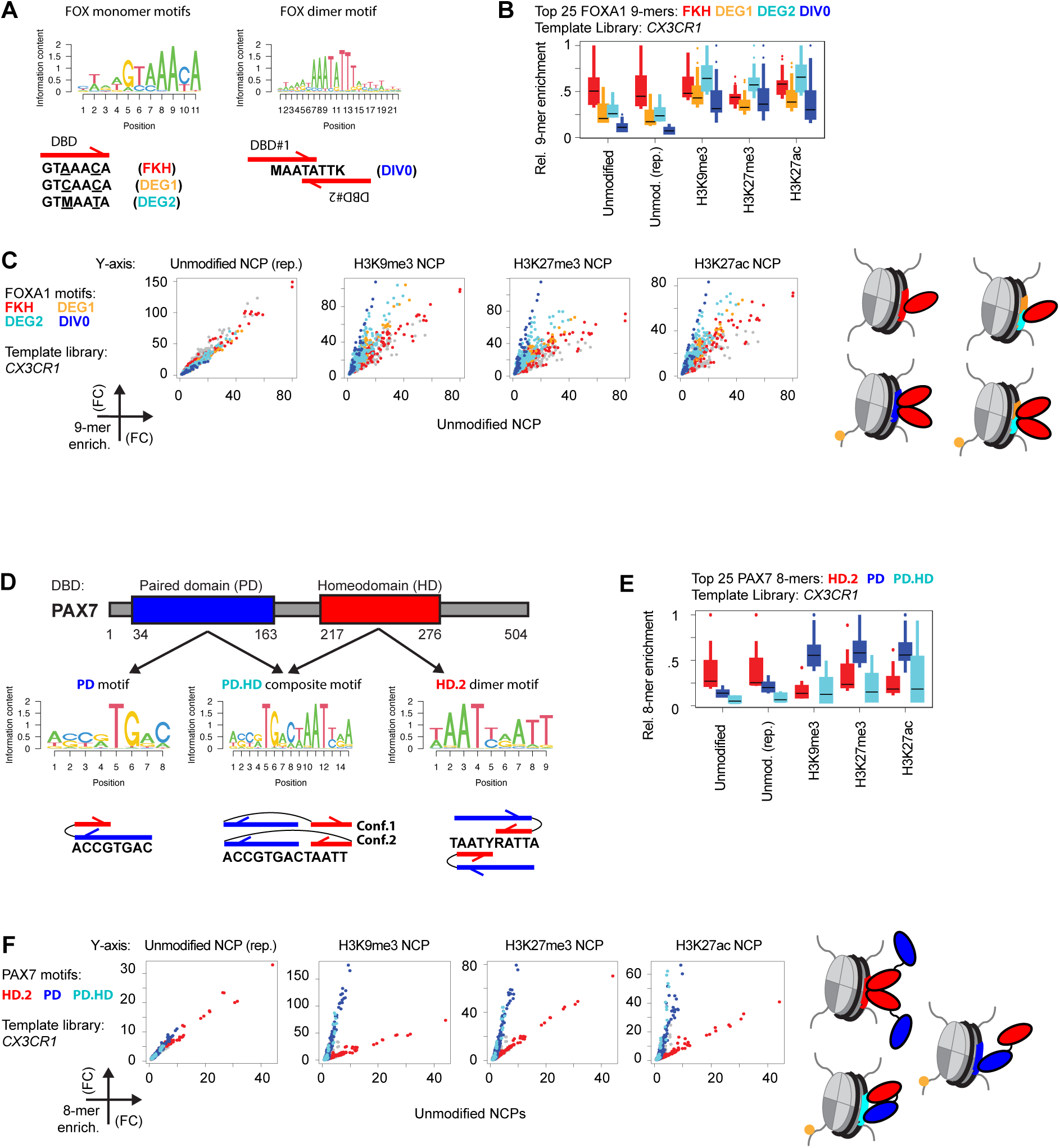
Epigenetic histone modifications modulate the NCP binding specificity of FOXA1 and PAX7. (**A**) Position weight matrix (PWM) motifs and consensus sequences *(bottom)* for the *in vivo* binding specificity of FOXA1, based on FOXA1 ChIP-seq (Cis-BP ID: M08419) *(left)* and DNase-seq data (*36*) *(right)*. Red arrows: motif recognition configuration (*i.e.*, monomer or dimer) of FOXA1 DBDs. M = C/A; K = T/G. (**B**, **C**) For the FOXA1-selected *CX3CR1*-based NCP assembled with the indicated histone 3, 9-mer enrichment values color-coded according to their match to the FOXA1 consensus sequences shown in (A). In (B), 9-mer enrichment values were calculated relative to the max 9-mer enrichment of the library. “(rep.)”: independent replicate PIONEAR-Seq assay using the NCP unmodified library. In (C), 9-mer enrichment values are calculated as in Fig. 1B. Gray dots in the scatterplots represent 9-mers that do not match any of the 4 FOXA1 consensus sequences shown in (A). Schemas on the right represent the favored FOXA1:NCP configuration for each consensus sequence. The same patterns appear in independent and template replicates, as in Fig. 1 (fig. S4). (**D**) Schematic organization of DBDs in human PAX7 (Uniprot ID P23759), their specificity (PWM motifs and consensus sequences) in ChIP-seq data (*41*) and domain configuration for DNA motif recognition. Red and blue arrows represent PD and HD DBDs, respectively, connected by the linker. Two alternative configurations (Conf.1 and Conf.2) can recognize PD.HD. Y = C/T; R = A/G. **(E**, **F)** Same as (B, C), but for the enrichment of 8-mers bound by PAX7 in *CX3CR1*-based NCP libraries. Boxes in the barplots and points in the scatterplots are color-coded according to their match to the PAX7 consensus sequences shown in (D). The same patterns appear in both independent and template replicates, as in Fig. 1 (fig. S5). In (F), the schemas on the right represent the favored PAX7:NCP configuration for binding to each recognition motif.

PAX7 is a member of the paired homeodomain family and is known to access and activate regulatory elements in heterochromatin in pituitary and muscle differentiation (*38*). PAX7 can assume multiple DNA binding modes using its two DBDs, a paired domain (PD) and a homeodomain (HD) (Fig. 2D). The PD contains two consecutive helix-turn-helix subdomains and recognizes the sequence GTCACGGT both alone (PD) (*39*) and as part of a composite motif (PD.HD) with the HD (*40, 41*) (Fig. 2D). Without the PD, the HD stably binds via DNA-mediated homodimerization to the HD.2 sequence TAATYRATTA (*42*) (Fig. 2D). In agreement with these prior observations, the PD, PD.HD and HD.2 motifs were enriched in PAX7 PIONEAR-seq data (Fig. 2E, F). However, while PAX7 favored the HD.2 motif on unmodified NCPs, it bound preferentially to the PD and PD.HD motifs on modified NCPs (Fig. 2E, F; fig. S5). Prior studies that profiled the genomic occupancy of PAX7 in pituitary cells found that it recognizes chromatin mostly through binding the PD.HD motif (*41*); our results suggest that it does so via modified nucleosomes. Altogether, our PIONEAR-seq data on modified NCPs show that the presence of histone PTMs can regulate the DNA binding specificity of both FOXA1 and PAX7, with the PTMs reconfiguring their preferences for binding nucleosomes as TF dimers rather than as monomers.

## H3K27 acetylation enhances TF preferences for binding to nucleosome ends

Although the PTMs we assayed by PIONEAR-Seq occur at different residues (H3K9, H3K27), add different chemical moieties (trimethylation, acetylation) and mark chromatin regions with different functions (constitutive or facultative repression, active), they caused similar changes in binding specificity for both FOXA1 (Fig. 2B, C) and PAX7 (Fig. 2E, F). We previously showed that the positional preference of TF binding to NCPs can be regulated independently of changes to sequence specificity (*13*). Therefore, we asked whether each PTM modulates the TF positional preference on NCPs differently. In PAX7 PIONEAR-seq data for unmodified and all three modified NCP libraries, all three PAX7 motifs (PD, PD.HD, HD.2) were enriched toward the left end of *CX3CR1*-based NCPs (Fig. 3A; fig. S6A) and towards the right end of *NRCAM*-based NCPs (fig. S6B), in agreement with our prior study showing that these ends are intrinsically more accessible for TF binding because of the flexibility profiles of these NCPs’ DNA sequences (*13*). In contrast, PAX7 did not show strong positional preference for binding on the ‘free DNA’ negative control (Fig. 3A; fig. S6). Enrichment of HD.2 8-mers (*i.e.*, AATCGATT) along *CX3CR1*-based NCPs was characterized by ∼10-bp periodicity on the left end of the template (*i.e.*, bp 34 and 45) (Fig. 3A), likely due to the phase of the DNA double helix as it wraps around the histone octamer, which may accommodate the PAX7 dimer in specific orientations that do not clash with the histone octamer.

**Fig. 3.**
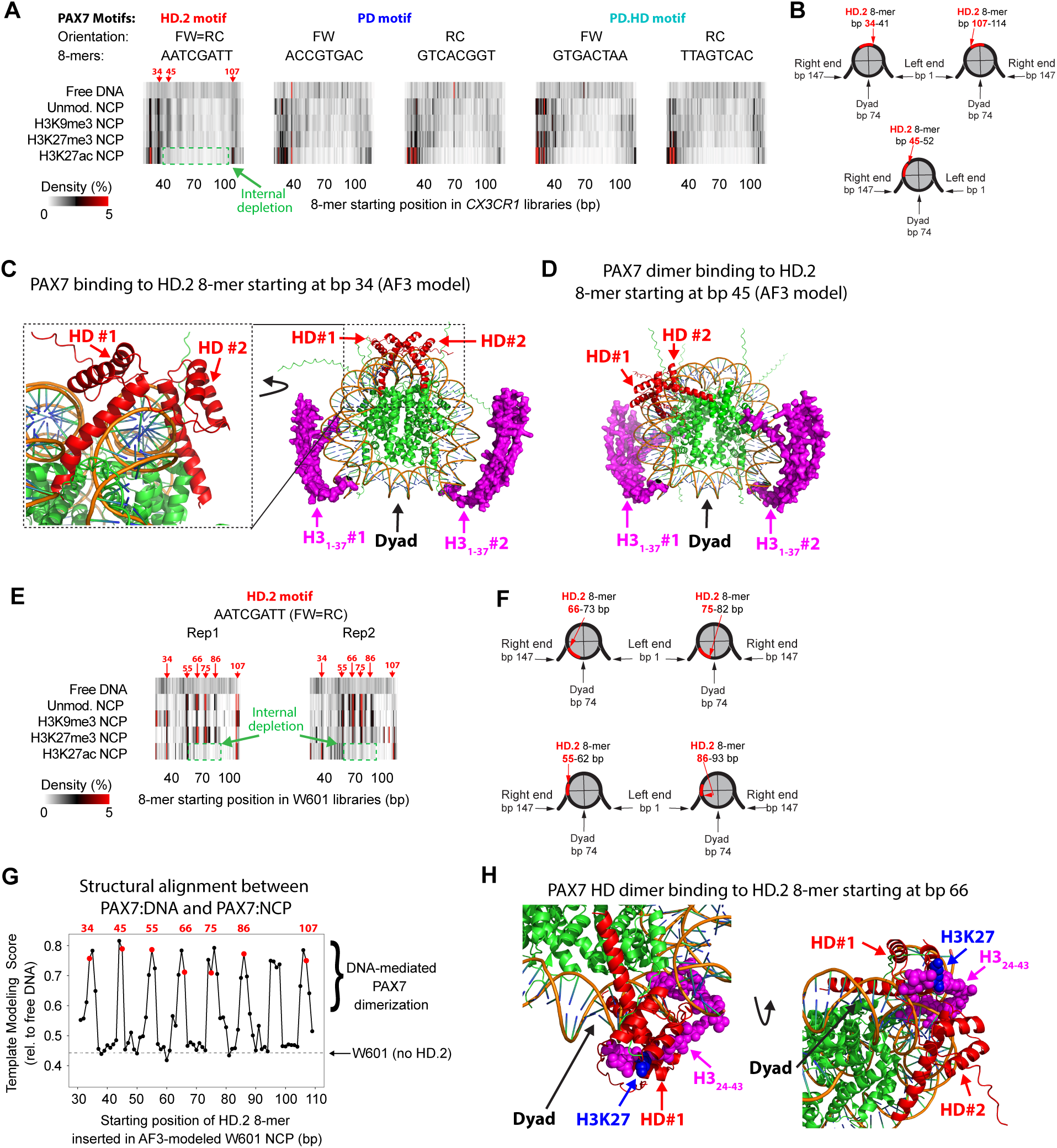
H3K27ac prevents PAX7 binding to NCP internal sites. **(A)** Density profiles of the indicated 8-mers along the N_20_ random tiles in PAX7-selected *CX3CR1* libraries assembled with different H3 tail PTMs (rows). Each 8-mer represents a PAX7 motif from Fig. 2D with a specific orientation. Green dashed boxes highlight internal depletion of PAX7 binding to H3K27ac NCPs, as compared to the other NCPs. Red arrows and numbers highlight preferred positions for PAX7 binding to HD.2 motifs in *CX3CR1*-based NCPs. The same patterns appear in independent replicate PIONEAR-Seq assays (fig. S6A). **(B)** For the density peaks at bp 34, 45 and 107 in (A), a schematic representation of the bound 8-mers (red) within the PAX7-bound NCPs. In the schema for the 8-mer at bp 107, the NCP is rotated 180 degrees about the vertical axis to highlight its symmetrical identity to the one at bp 34. Red arrows point to the starting bp of the 8-mers (red segments). (**C**) AlphaFold3 model of a complex with two PAX7 HD DBDs and an NCP with the HD.2 8-mer at bp 34. Left inset: rotated zoom-in to highlight the DNA-mediated dimerization of PAX7 on the NCP. The H3 tails (a.a. 1-37) of 6 AF3 structural seeds aligned to the presented structure are shown. Red: PAX7 HDs; orange: DNA backbone; green: histones, except for the H3 N-terminal tail (magenta and displayed as 3D-contour of solvent-accessible surface). (**D**) As in (C), but for an HD.2 8-mer inserted at bp 45. (**E**) As in the left heatmap of (A), but for W601-based NCPs. Rep1, Rep2: results from two independent replicate PIONEAR-Seq assays. Red arrows and numbers: preferred positions for PAX7 binding to the HD.2 motif. (**F**) As in (B), but for the indicated HD.2 8-mer positions shown in (E). (**G**) Template modeling (TM) scores between an AF3-generated reference model of PAX7-bound HD.2 as free DNA and a series of AF3-modeled PAX7-bound NCPs carrying HD.2 at the indicated positions (see Methods). Each structure included 2 copies of the PAX7 HD.2. For each HD.2 position, the TM scores of 5 AF3 structures were calculated and their max value was depicted (black points). Red points: HD.2 positions of positions highlighted in (A) and (E). The gray dashed line corresponds to the alignment of the W601 NCP without HD.2 insertion. (**H**) Zoom-ins of the structural alignment between an NCP that contains a cryo-EM-resolved H3 tail (PDB: 8G8G) and 2 PAX7 HD DBDs predicted by AF3 to recognize their cognate 8-mer starting at bp 66 of the NCP. Coloring as in (C), except that H3K27 is highlighted in blue.

In contrast, H3K27ac – but neither H3K27me3 nor N3K9me3 – resulted in a depletion of PAX7 binding to the most internal regions of the nucleosomes (Fig. 3A). In the case of the HD.2 8-mers in *CX3CR1*-based NCPs, this H3K27ac-specific depletion resulted in loss of binding at bp 45 and gain of binding at bp 107, which is symmetrical, relative to the dyad at bp 74, to the gain at bp 34 (Fig. 3A, B). To corroborate that these sites represent sterically permitted PAX7 binding along nucleosomes, we used AlphaFold3 (AF3) (*43*) to model two PAX7 DBDs bound to HD.2 8-mers inserted within an unmodified NCP. Indeed, AF3 models confirmed that these positions (*i.e.*, bp 34 and 45) can accommodate both PAX7 DBDs without causing steric hindrance with the structured domains of the histones (Fig. 3C), while sites located at nearby positions (*i.e.*, bp 32, 36, 43, 47) do not allow such DNA-mediated PAX7 dimerization (fig. S7). Intriguingly, the AF3 models indicate that the H3 tails’ range of conformations reaches PAX7 DBDs bound at bp 45 (Fig. 3D) with higher probability than for PAX7 bound at bp 34 (Fig. 3C), suggesting that the loss of signal at bp 45 in the presence of H3K27ac (left heatmap, Fig. 3A) may be due to a direct, inhibitory interaction between the H3 tail and PAX7.

To gain further support for our observation that H3K27ac enhances PAX7 binding towards the NCPs’ DNA ends, we examined PAX7 positional preference for HD.2 in W601-based NCPs (Fig. 3E), which we previously found to show preferential TF binding near the dyad of unmodified NCPs (*13*). Indeed, W601-based NCPs bound by PAX7 exhibited 4 preferential sites of binding (*i.e.*, bp 55, 66, 75, 86) to HD.2 8-mers near the dyad (Fig. 3E). Similar to the peaks at bp 34 and 107 (Fig. 3A), these 8-mer positions represent two symmetrical configurations (on the left (bp 55, 66) and on the right (bp 75, 86) of the dyad (Fig. 3F)) that sterically allow DNA-mediated PAX7 dimerization on the NCPs according to AF3 models (fig. S7).

To systematically scan for DNA-mediated PAX7 dimerization on the NCPs, we generated AF3 models for the binding of 2 PAX7 HDs to NCPs carrying the HD.2 8-mers at each internal position from bp 31 to 110, encompassing all the preferential sites of PAX7 binding detected by PIONEAR-seq (Fig. 3A, E). We also generated an AF3 model for 2 PAX7 HDs bound to HD.2 on free DNA (fig. S8). We performed a structural alignment of this reference dimer structure to each NCP model using US-align (*44*). We then obtained Template Modeling (TM) score to quantify the capacity of the AF3-modeled nucleosome structure to accommodate a PAX7 HD dimer at each position. The resulting TM score profile along the NCP showed that the positions enriched for HD.2 8-mers in PAX7 PIONEAR-seq data (bp 34, 45, 55, 66, 75, 86, 107) specifically allow for binding by PAX7 dimers to the NCPs (Fig. 3G). In agreement with our PIONEAR-seq data for PAX7 binding to *CX3CR1*-based NCPs (Fig. 3A), the presence of H3K27ac also inhibited binding of PAX7 to HD.2 8-mers at internal positions of W601-based NCPs, while favoring binding at bp 34 and 107 (Fig. 3E). Since AF3 does not provide information about conformational dynamics, and intrinsically disordered histone tails are particularly difficult to interpret from AF3-modeled structures, we superimposed a cryo-EM structure of a resolved H3 tail (residues 24-43) (*23*) onto the AF3 predicted PAX7-bound NCP to gain insights on the potential locations of the H3 tail relative to NCP-bound PAX7. The results showed that H3K27 likely interacts with PAX7 bound to internal NCP positions (*i.e.*, bp 66) (Fig. 3H), suggesting that H3K27ac might directly inhibit the interactions between PAX7 and H3.

On W601-based unmodified NCPs, FOXA1 bound to the DIV0 motif preferentially at specific internal positions (Fig. 4A; fig. S9A). Consistent with our observations for PAX7 (Fig. 3E), the presence of H3K27ac (but not H3K9me3 or H3K27me3) inhibited FOXA1 binding to DIV0 sites at internal positions (*i.e.*, bp 70-71), while enhancing binding to DIV0 near the NCP ends (*i.e.*, bp 29, 101) (Fig. 4B; fig. S9B, C).

**Fig. 4.**
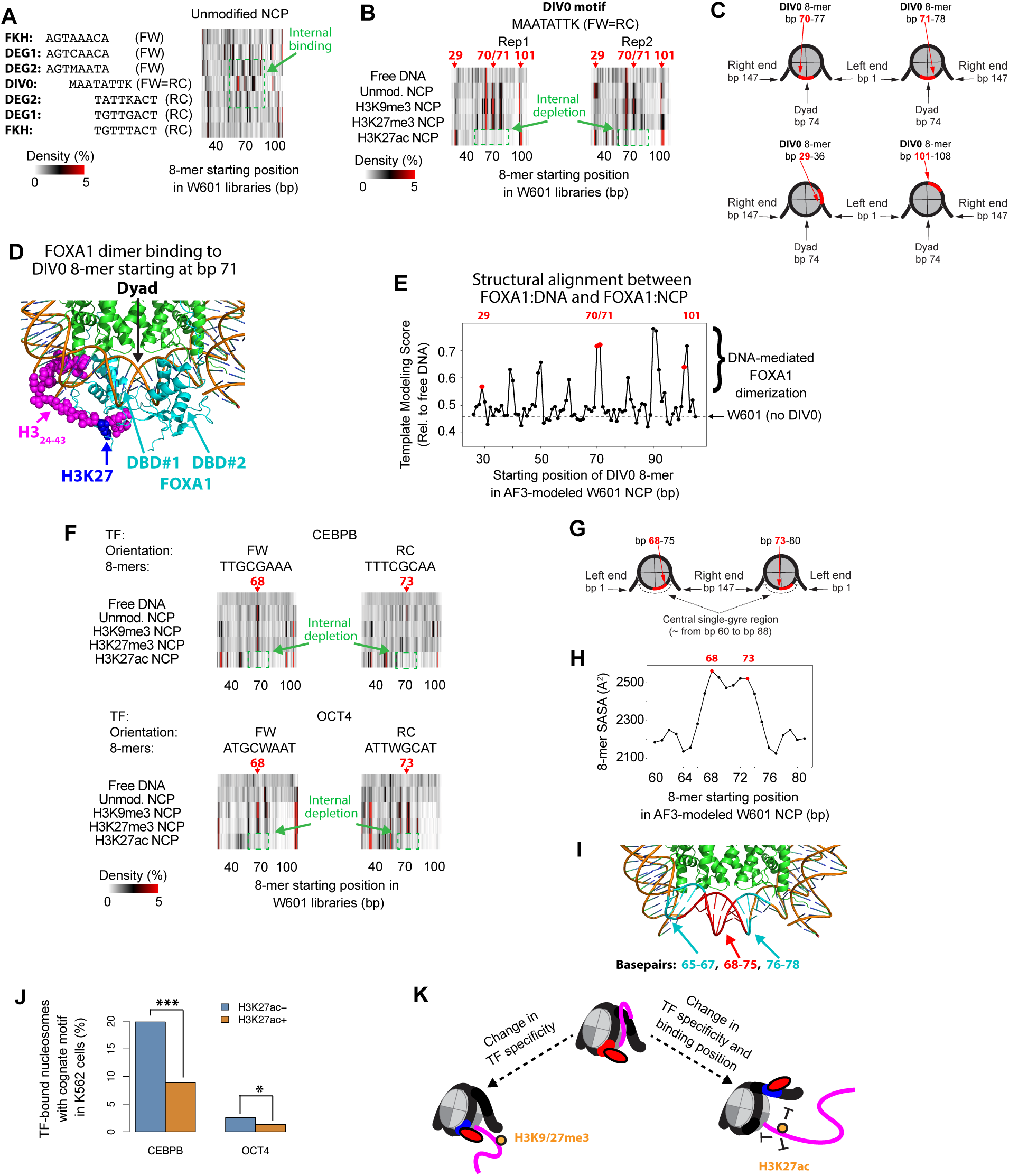
H3K27ac enhances TF end-binding mode. **(A)** Density profiles of the indicated 8-mers along the N_20_ random tiles bound by FOXA1 in unmodified W601 NCP libraries. Each 8-mer is representative of one FOXA1 consensus sequence from Fig. 2A and is presented in both forward (FW) and reverse complementary (RC) orientation. Green dashed box highlights the binding of FOXA1 to the DIV0 and DEG2 8-mers preferentially at or near the dyad. M = C/A; K = T/G. **(B)** Two independent replicates for the density profiles of the DIV0 8-mer along the N_20_ random tiles for FOXA1-selected W601 NCP libraries assembled with different H3 tail PTMs (rows). Red arrows and numbers highlight preferential 8-mer positions for FOXA1 binding to W601 NCPs. Green dashed box highlights depletion of FOXA1 binding at or near the dyad of H3K27ac NCPs. **(C)** Schematic representations of the start and end positions of the indicated DIV0 8-mer insertions within 147-bp nucleosomal DNA. Red numbers: 8-mer starting bp position. For DIV0 positions at bp 71 and 101, the NCP was rotated 180 degrees about the vertical axis to highlight its symmetrical identity to bp 70 and 29, respectively. Red arrows point to the starting bp position of the 8-mers (red segments). **(D)** Zoom-in of the structural alignment between an NCP that contains a cryo-EM-resolved H3 tail (PDB: 8G8G) and 2 FOXA1 DBDs predicted by AF3 to recognize the DIV0 8-mer starting at bp 71. Colors as in Fig. 3H, expect here for FOXA1 instead of PAX7. **(E)** TM scores quantifying the alignment between an AF3 model of FOXA1 bound to DIV0 on free DNA and a series of AF3-modeled FOXA1-bound NCPs carrying DIV0 at the indicated positions. Each structure included 2 copies of the FOXA1 DBD. Black points: as in Fig. 3G. Red points: DIV0 positions highlighted in (B). The gray dashed line corresponds to the alignment of the W601 NCP without the DIV0 insertion. **(F)** Positional density profiles of the indicated 8-mers along the N_20_ random tiles for W601 libraries selected by CEBPB *(above)* and OCT4 *(below)* in different NCP types (rows). Each 8-mer represents the consensus sequence of the associated TF and is presented in both forward (FW) and reverse complementary (RC) orientations. Red arrows and numbers highlight the preferred positions for CEBPB and OCT4 binding to W601 NCPs. Green dashed boxes: as in (B) but for CEBPB *(above)* and OCT4 *(below)*. **(G)** Schematic representation of the 8-mer positions at bp 68 and bp 73 (red) within a 147-bp NCP. For 8-mers at bp 73, the NCP was rotated 180 degrees about the vertical axis to highlight their symmetrical configurations with the 8-mer at bp 68. Red arrows point to the starting position of the 8-mer. Dashed arcs indicate the single-gyre region. **(H)** Solvent-accessible Surface Area (SASA) calculated by AlphaFold3 for 8-mers starting at the indicated bp position within the W601 template in an AF3-modeled NCP. 8-mers starting at positions bp 61 to bp 79 represent 8-mers located completely within the single-gyre DNA region (bp 60-88) near the dyad (bp 74). Local SASA maxima (bp 68 and 73) are highlighted in red. **(I)** Zoom-in of the NCP structure (PDB: 8G8G) around the dyad. DNA base pairs are color-coded to highlight the 8-mer starting at bp 68 (red) and its flanking regions (cyan). **(J)** Percentages of TF-associated nucleosomes containing either CEBPB *(left)* or OCT4 (*right)* binding motifs. Significant differences (*** *P* < 0.001, * *P* < 0.05, Fisher’s exact test) in corresponding motif frequencies in H3K27ac+ vs. H3K27ac-nucleosomes. **(K)** Schematic of a model for the molecular interactions between TFs and NCPs in their unmodified state or modified with the indicated PTMs.

AF3 models of FOXA1 binding to NCPs with a DIV0 site at these positions, which are located almost symmetrically on the left (bp 29 and 70) and right (bp 71 and 101) of the dyad (Fig. 4C), showed sterically favorable configurations for FOXA1 dimer binding on the NCP, in contrast to nearby positions (*i.e.*, bp 69 and 72) (Fig. 4D; fig. S10A). As seen for PAX7 (Fig. 3H), binding to these preferred internal positions brings FOXA1 DBDs in the vicinity of the H3 tail (Fig. 4D), potentially enabling interactions regulated by the acetylation of H3K27. Positions of preferred FOXA1 binding as observed in PIONEAR-seq data had high TM scores in an alignment of a free DNA reference model of DIV0-bound FOXA1 dimer (fig. S10B) to FOXA1 DBD binding to DIV0 sites tiling the NCP (from bp 26 to 105) (Fig. 4E). Altogether, these results support our model of DNA-mediated FOXA1 dimerization on NCPs, as also seen for PAX7 (Fig. 3G).

To determine whether H3K27ac might inhibit binding by a broader set of TFs to internal NCP regions, we analyzed PIONEAR-Seq data for additional TFs. We previously showed that CEBPB preferentially binds its cognate 8-mer TTGCGAAA near the dyad of W601 NCPs at bp 68 and 73 in the forward and reverse complementary orientations, respectively (*13*). Here, we found that H3K27ac inhibited internal binding of CEBPB near the dyad, with CEBPB instead binding preferentially near the NCP ends (Fig. 4F; fig. S11). Similarly, H3K27ac inhibited binding of OCT4 to its cognate 8-mer ATGCAAAT (Fig. 4F; fig. S11), and of FOXA1 to the DEG2 8-mer (fig. S9C), at bp 68 and 73. Altogether, these results indicate that while these 8-mer positions, which are located fully within the NCP’s central single-gyre region where inter-gyre steric obstruction is absent (Fig. 4G), are intrinsically favorable for TF binding, in agreement with their estimated solvent-accessible surface area (Fig. 4H, I), H3K27ac nevertheless inhibits TF binding at these positions.

To determine whether these inhibitory effects of H3K27ac occur *in vivo*, we analyzed ENCODE ChIP-Seq and MNase-Seq data to identify nucleosomes marked by H3K27ac and bound by CEBPB and OCT4 in K562 cells (*45*). 23,861of the 61,637 nucleosomes that overlapped CEBPB ChIP-seq peaks also overlapped H3K27ac ChIP-seq peaks (H3K27ac+ nucleosomes). Only 8.9% of the H3K27ac+ nucleosomes contained a CEBPB binding site motif, in contrast to 19.9% of the H3K27ac-nucleosomes (Fig. 4J), indicating that CEBPB direct binding to nucleosomes is significantly depleted for binding to H3K27ac+ nucleosomes (*p* < 2.2 x 10^-16^, Fisher’s exact test). Similarly, OCT4 direct binding to nucleosomes was significantly depleted among H3K27ac+ nucleosomes (*p* = 0.017, Fisher’s exact test). Together with the PIONEAR-seq data, these results suggest that H3K27ac inhibits nucleosome binding by these TFs *in vivo*.

Our results reveal that PTMs on nucleosomes can directly modulate both the DNA sequence specificity and positional preference of TF binding to nucleosomes (Fig. 4K) in the absence of chromatin readers, thus revealing an additional *cis*-regulatory layer in the epigenome. Consistent with recent structural studies that showed that TFs interact with the histone tails of nucleosomes (*23, 46*), here we report PTM-dependent changes of binding specificity for PAX7 and FOXA1 (Fig. 2). Similar binding specificity changes being induced by H3K27ac, H3K9me3 and H3K27me3 (Fig. 2C, F) suggests that these PTMs modify TF motif recognition by similarly modulating histone tail interactions and/or dynamics. H3K27ac, but not H3K27me3, inducing changes in the preferential positions of TF binding within nucleosomes (Fig. 3, 4) is consistent with prior evidence that histone tail acetylation, which neutralizes the positive charge of lysines, increases the DNA unwrapping of the nearby nucleosome ends (*17-20*) more effectively than does lysine methylation (*21*); our results suggests that such unwrapping might enable increased TF binding at nucleosome ends. Future studies are needed to determine whether these TFs contain cryptic PTM-reader domains, as reported for other chromatin factors (*47*). PIONEAR-seq analysis of a broader set of TFs and epigenomic marks likely will provide further insights into the molecular mechanisms regulating TF nucleosome binding during cell fate transitions.

## Supporting information

Tables S1 through S5

## Acknowledgments

We thank Kwangwoon Lee, Mingxuan Wu and Phil Cole for technical assistance in the production of octamers and nucleosomes in the initial stages of this project. We thank members of the Bulyk lab for helpful discussion. We thank Steve Gisselbrecht, Kwangwoon Lee and Phil Cole for critical reading of the manuscript.

## Author contributions

MLB conceived this study. LM and MLB designed the experiments and analyses. LM performed all experiments and analyses. MLB supervised the research. LM prepared the figures. LM and MLB wrote the manuscript. Both authors approved the final manuscript.

## Data, code, and materials availability

All plasmids used in this study for *in vitro* production of proteins are available from the lead contact without restriction. Sequencing data have been deposited in GEO under the accession number GSE333423 and will be made publicly available upon acceptance of this manuscript for publication. Sequences used for AlphaFold3 structure predictions along with AlphaFold3 output files have been deposited in the Harvard Dataverse at: https://doi.org/10.7910/DVN/RMT39Y.

## Supplementary Materials

Materials and Methods

Supplementary references

Figs. S1 to S11

Tables S1 to S5

