## Supplementary figures and images for "Histone modifications directly modulate transcription factor binding to nucleosomes"

### Figure S1

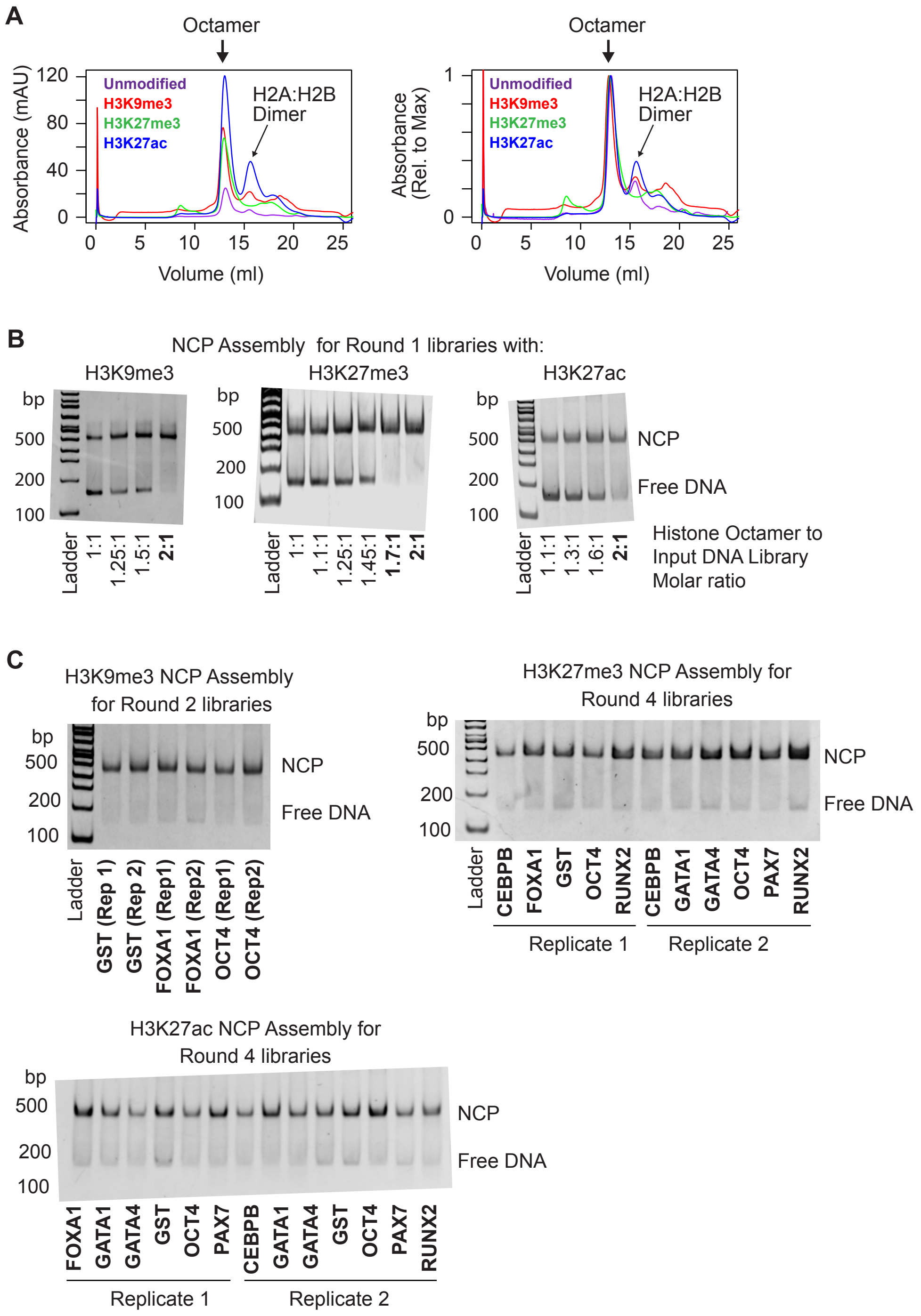

### Figure S2

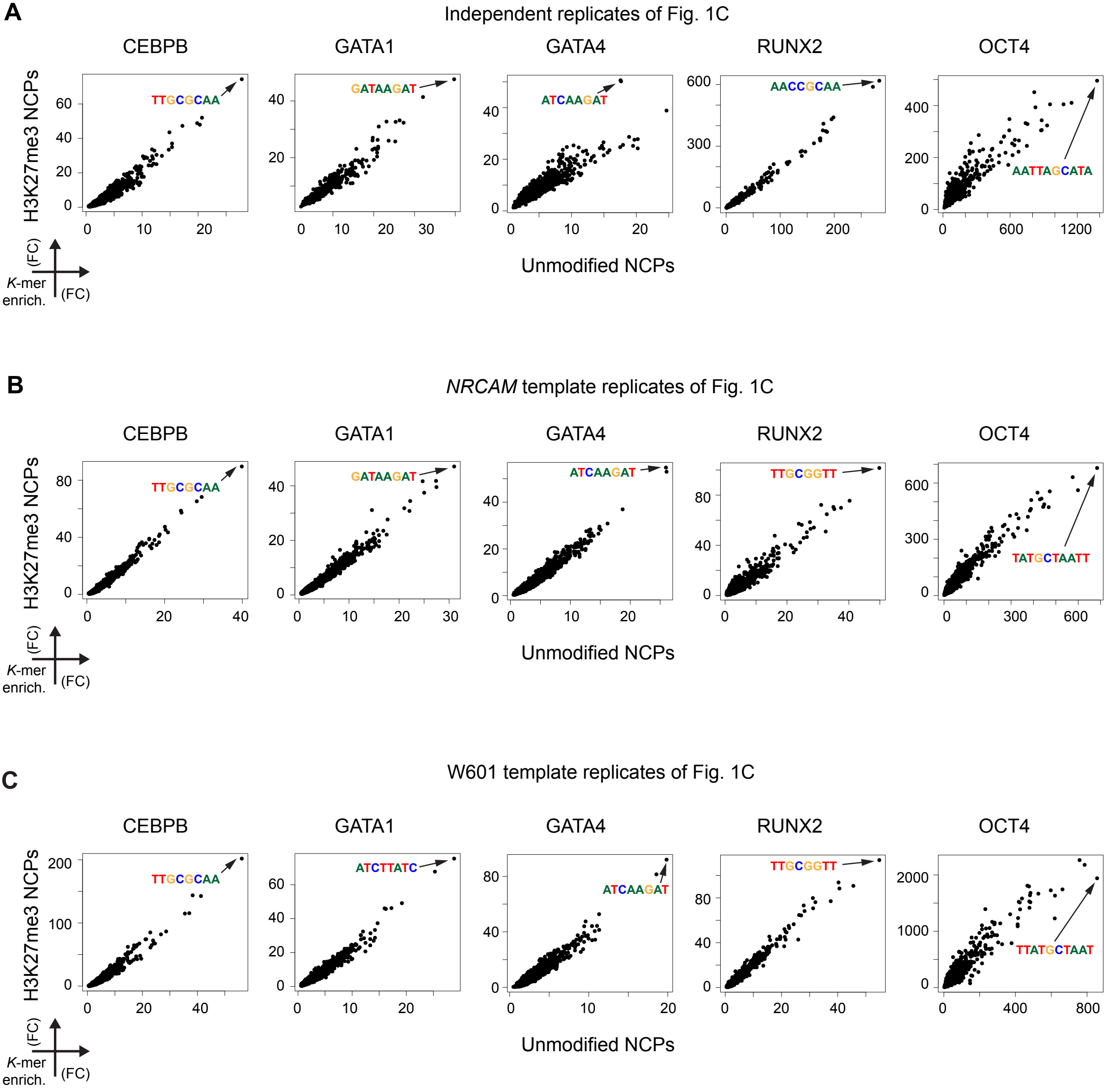

### Figure S3

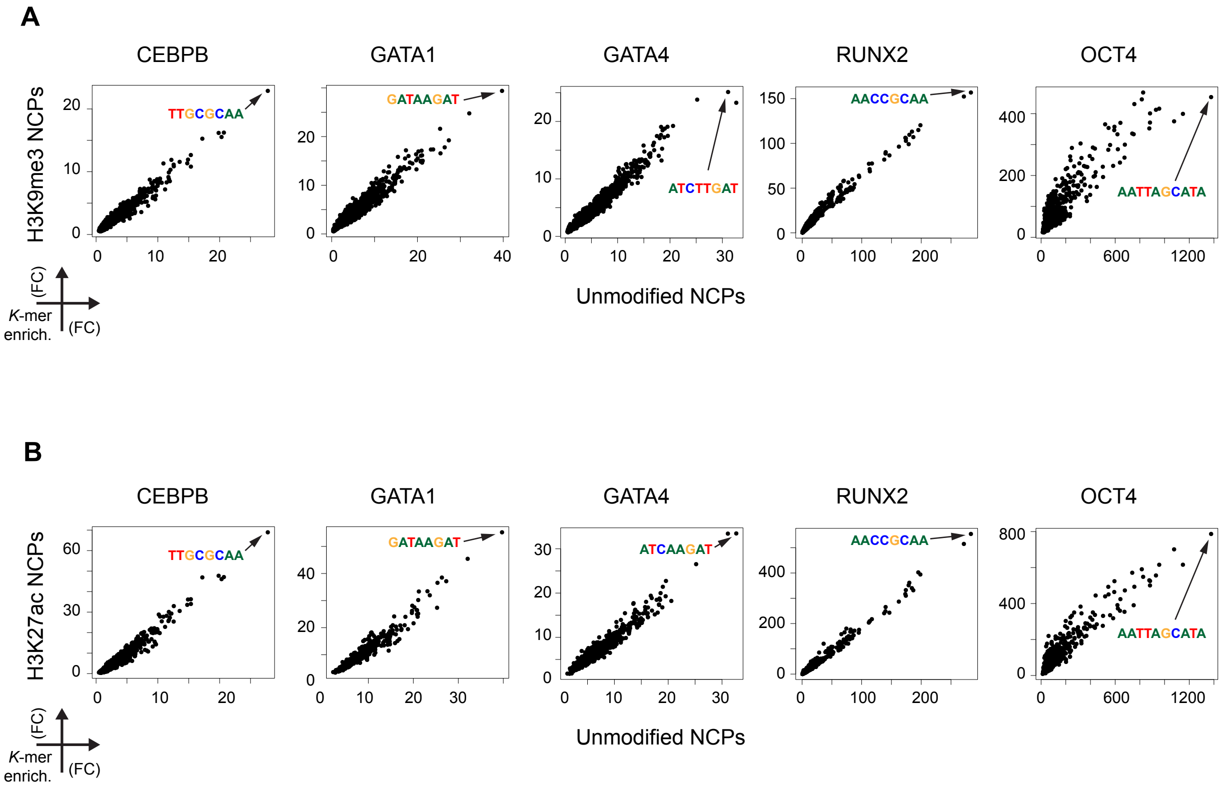

### Figure S4

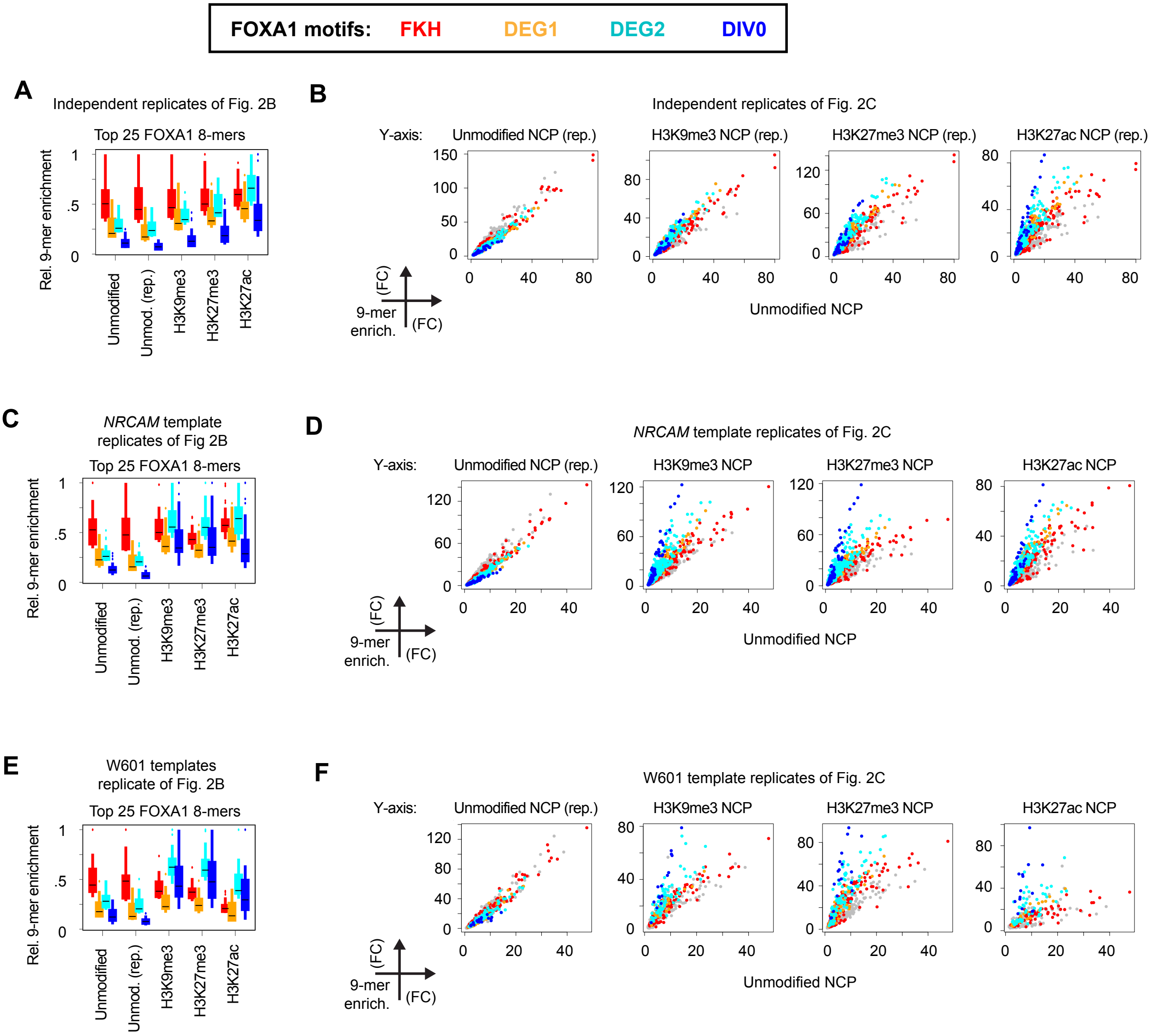

### Figure S5

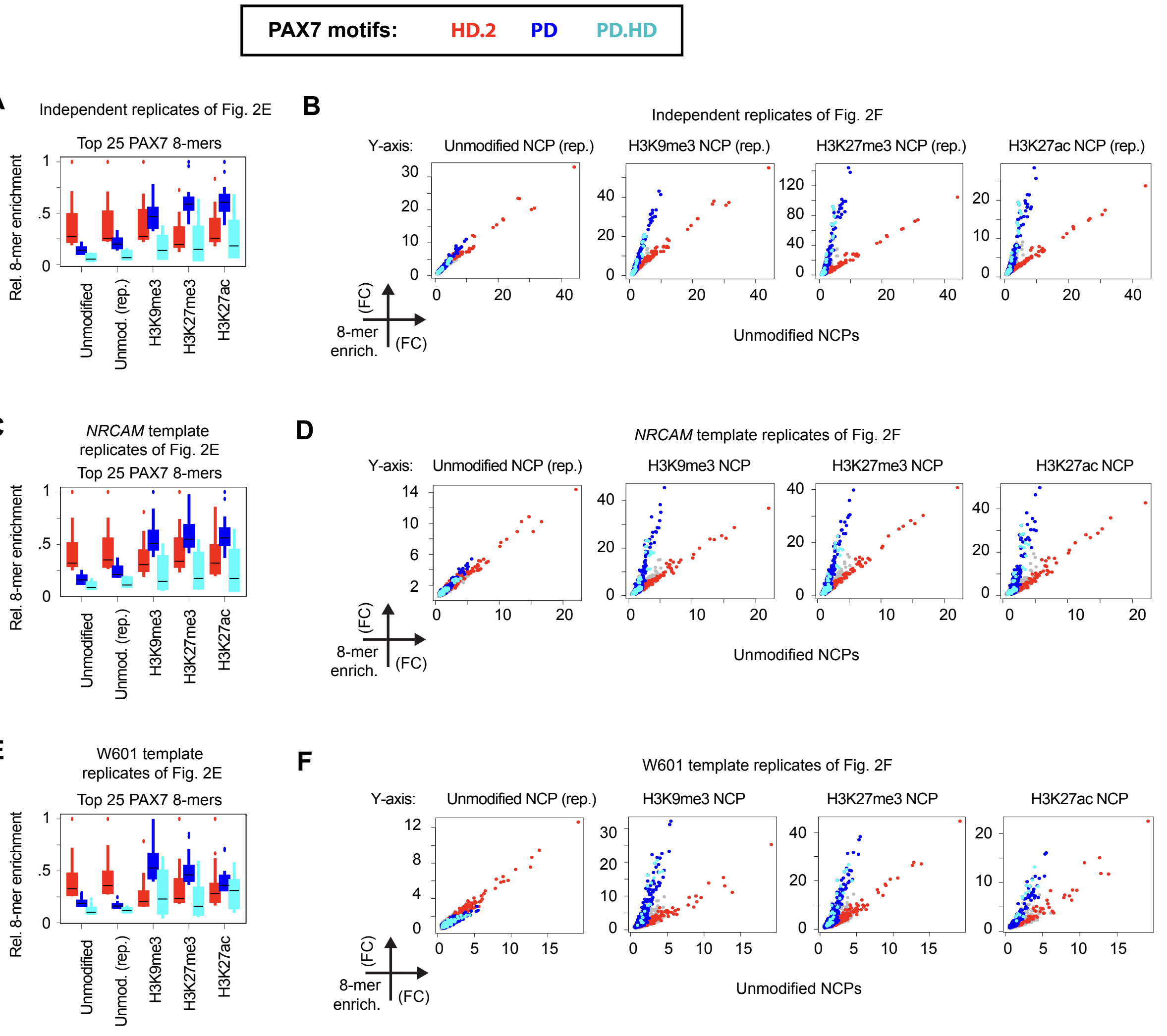

### Figure S6

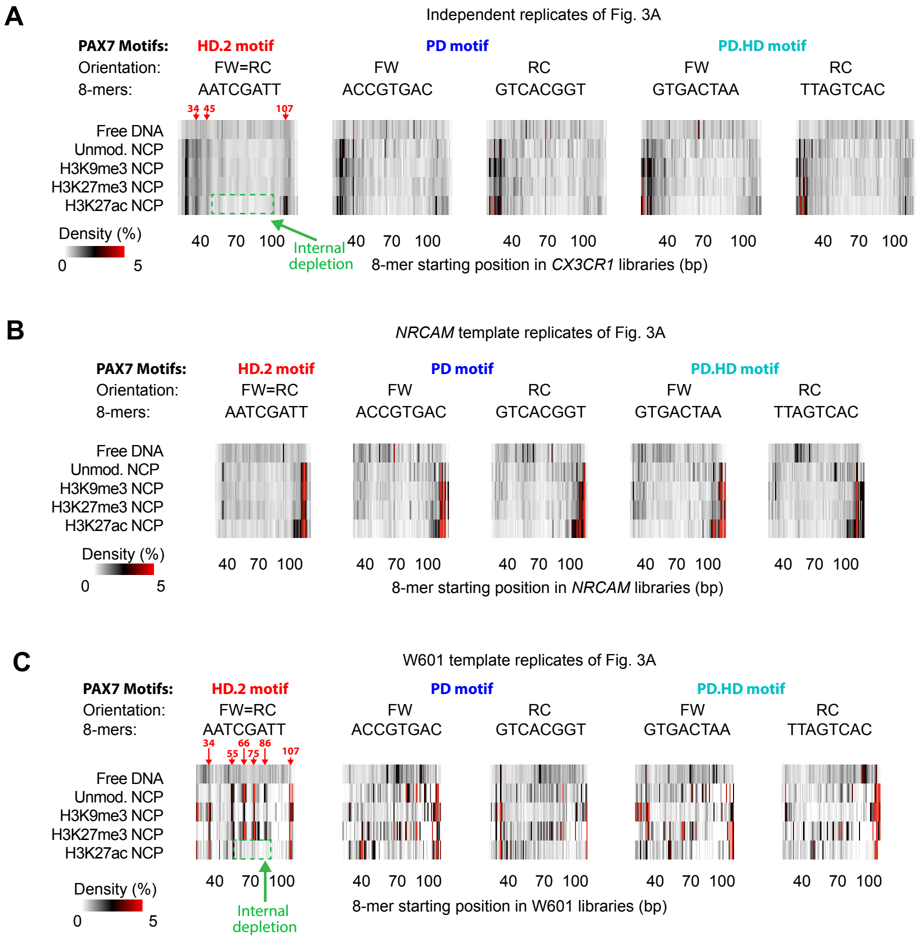

### Figure S7

HD.2 DNA 8-mer (AATCGATT) starting at :

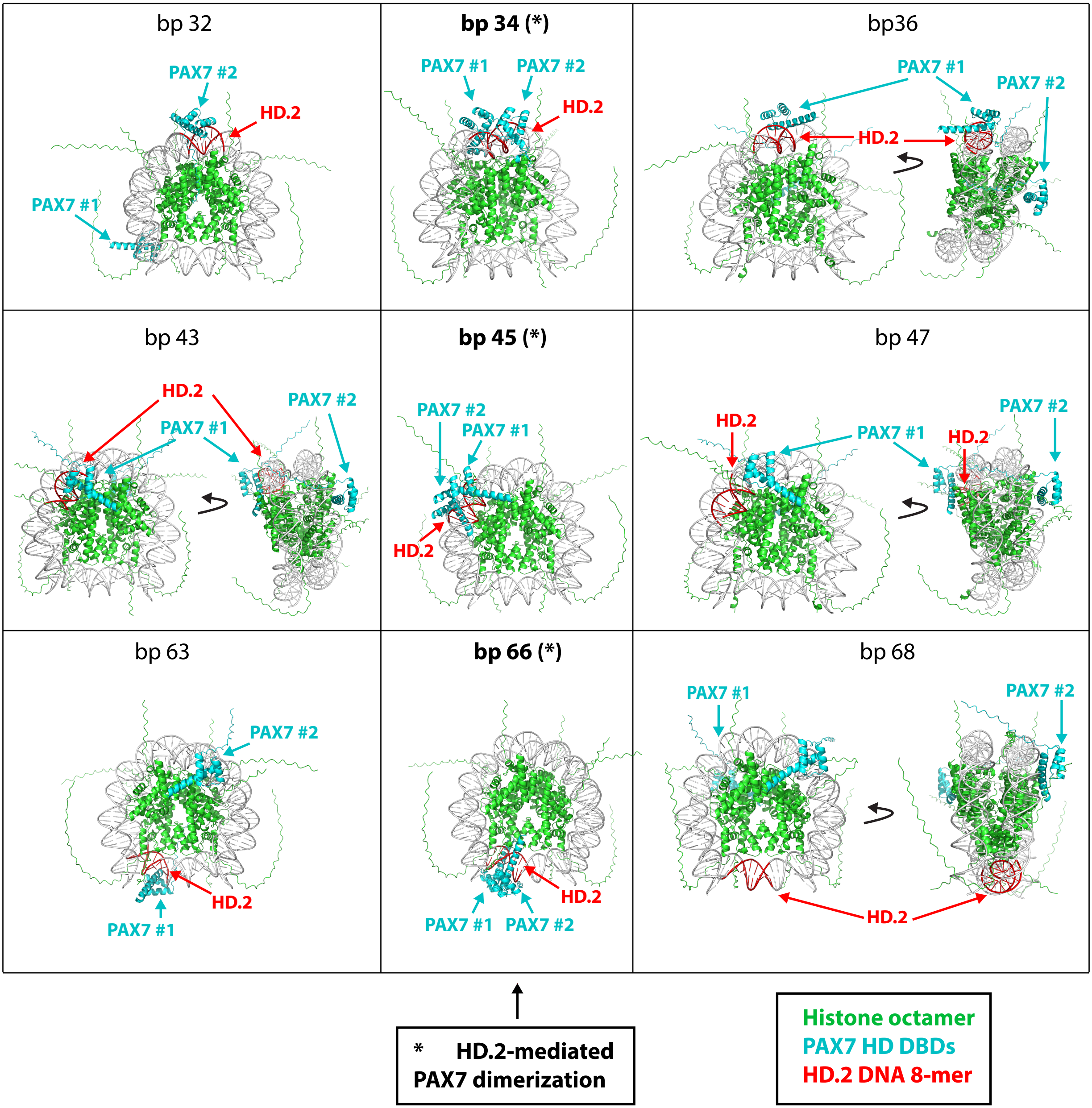


**Figure S7**

### Figure S8

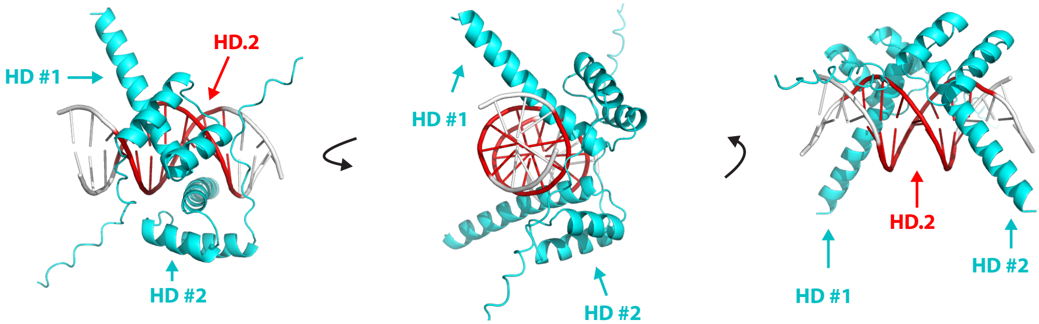


PAX7 HD DBDs HD.2 DNA 8-mer ( TATAATCGATTATA )

**Figure S8**

### Figure S9

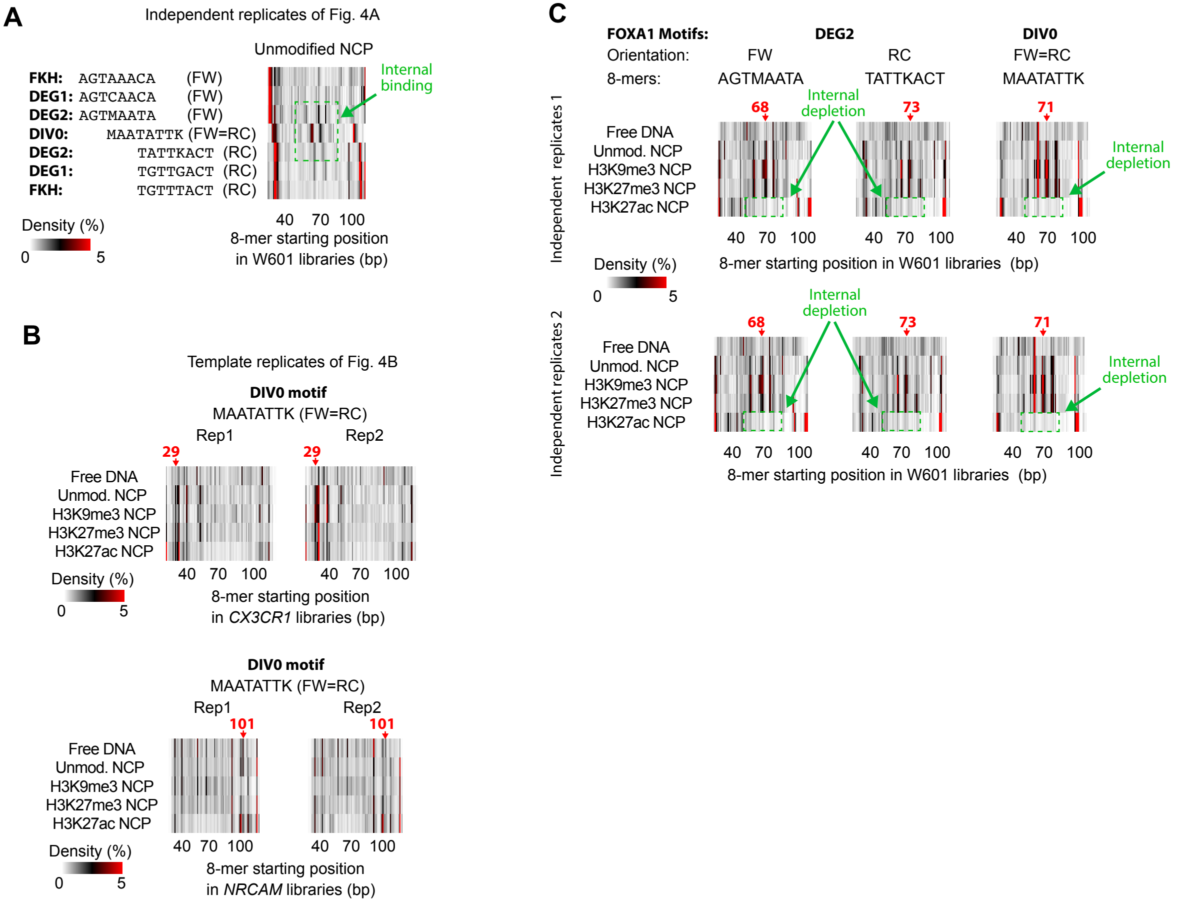

### Figure S10

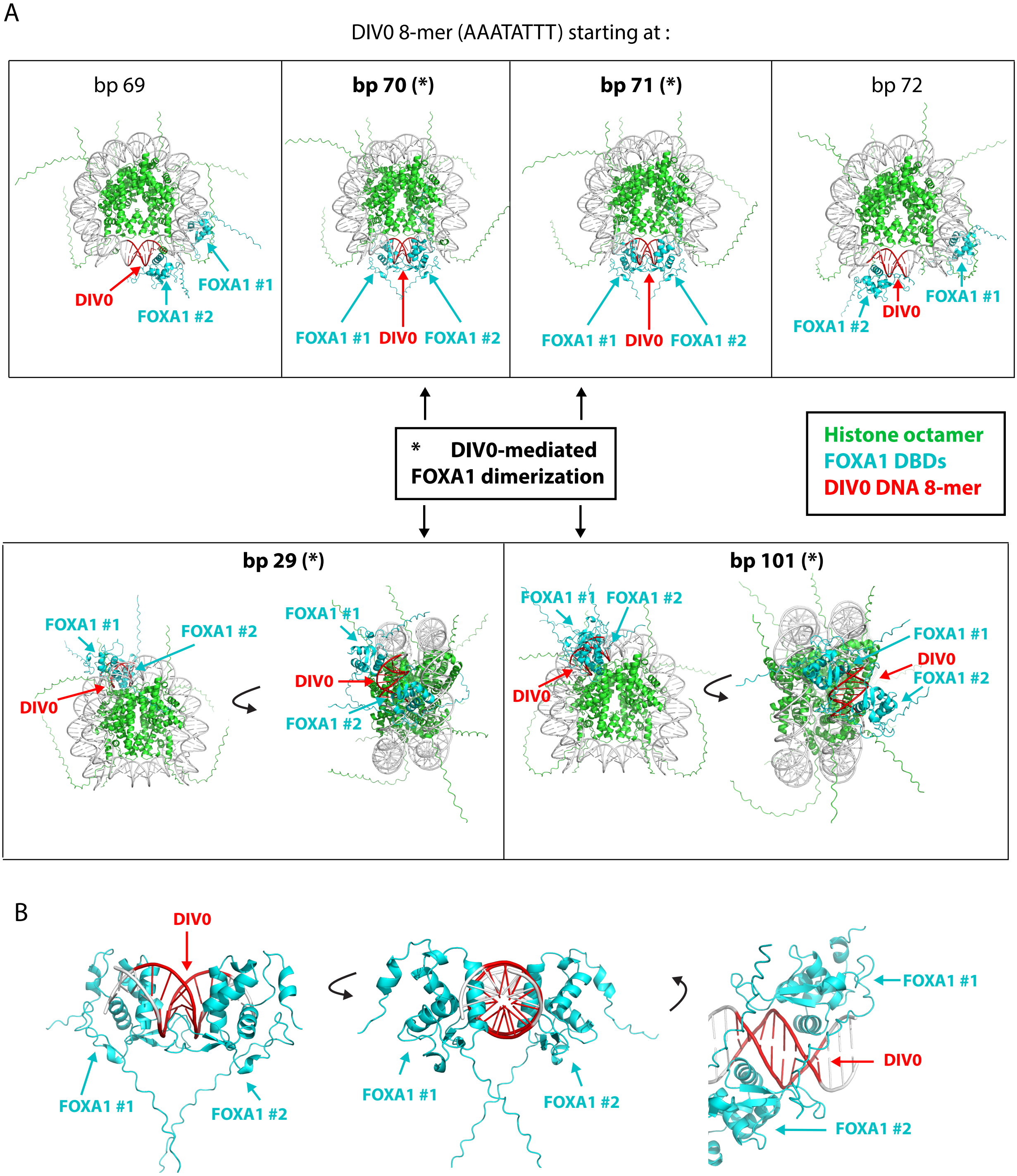

### Figure S11

Independent replicates of Fig. 4F

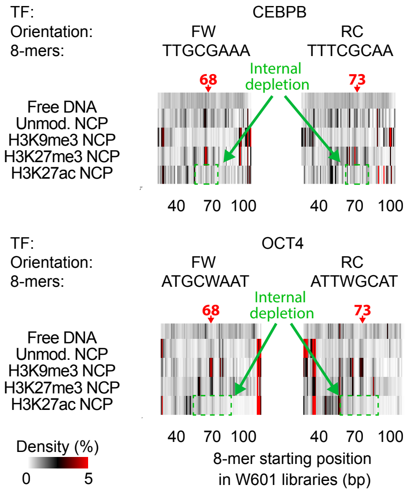


**Figure S11**
