## Supplementary material for "Histone modifications directly modulate transcription factor binding to nucleosomes": Suppl Materials text file

**This file includes:**

Materials and Methods
Legends for Figs. S1 to S11
Supplementary references

**Other Supplementary Materials include:**

Figs. S1 to S11
Tables S1 to S5

### Materials and Methods

PIONEAR-seq assay

PIONEAR-seq assays for nucleosomes that contained histone modifications were performed essentially as described previously for unmodified nucleosome (1), as summarized below and schematically depicted in Fig 1A. Briefly, DNA sequences that favor nucleosome positioning ('147-bp templates') are tiled with 20-bp random regions ('$N_{20}$') in the internal regions between fixed primers ('tiling'), which is then double-stranded, pooled together to form the 'Input DNA library' (Round 0) and assembled into nucleosome core particles (NCPs) using octamers reconstituted with histones of interest, such as unmodified H3 or H3 with PTMs ('NCP library assembly'). NCPs are incubated with GST-tagged human full-length transcription factors (TFs) to induce DNA specific binding and then pulled down using GST-specific beads ('TF-NCP binding and pull-down'). After 3 additional rounds of selection, the DNA libraries from Rounds 3 and 4 are sequenced.

For the assembly and purification of histone octamers, purified recombinant *H. sapiens* canonical histones were used. Unmodified (SKU ID: HH2A, HH2B, HH3 and HH4) as well as H3 histones that carry genetically encoded acetyl-lysine K27ac (2) (SKU: XH_H3K27AC) were purchased from Histone Source, aliquoted, lyophilized, and stored at -20 °C prior to use. Purified recombinant human H3 histones that carry trimethylation modifications as methyl-lysine analogs (3) were purchased from Active Motif (H3K9me3 (#31701), H3K27me3 (#31979)), aliquoted and stored at -80 °C prior to use. All histones were resuspended in unfolding buffer (6 M guanidine hydrochloride, 5 mM DTT and 20 mM Tris-Cl pH 7.5); the lyophilized histones were resuspended to a final concentration of 2-3 mg/ml, whereas the ones in suspension were brought to a final concentration of 0.2-0.3 mg/ml. H2A, H2B, H4 and one H3 type (unmodified H3, H3K9me3, H3K27me3 or H3K27ac) were then combined at a molar ratio of 1.1:1.1:1:1. The combined sample was incubated at room temperature for 30 minutes then dialyzed overnight at 4 °C twice against 900 ml refolding buffer (10 mM Tris-Cl pH 7.5, 2 M NaCl, 1 mM EDTA and 5 mM BME) and then purified by size exclusion chromatography (Fig. S1A). The FPLC fractions containing the octamer were pooled, concentrated to about 1.5 mg/ml and stored at 4 °C for up to 2 weeks.

For TF protein production, previously generated pDEST15 vectors containing codon-optimized sequences (Table S3) of the 7 TFs (CEBPB, FOXA1, GATA1, GATA4, OCT4/POU5F1, PAX7 and RUNX2) were used (*1*). These vectors, which add an N-terminal GST tag on the TFs, were verified by Sanger sequencing, and then used to express the tagged TFs using the NEB PURExpress kit (30 °C for 4 hours) were verified previously for yields and correct protein expression by Western blots (*1*). For each PIONEAR-seq experimental round, TF expression was performed fresh.

For the construction of PIONEAR-seq input DNA, two naturally occurring genomic 147-bp sequences (*CX3CR1*, *NRCAM*) (*4*) and one high-affinity synthetic 147-bp sequence (Widom 601) (*5*) were used as templates for their recognized ability to position nucleosomes (Table S1). The ~20-bp ends of each template were kept constant and used as primers, to ensure specific and efficient DNA amplification (Table S1). Within each template, multiple 20-bp internal segments were replaced with random sequences (‘tiles’) (Table S2). These tiles were shifted at 3-bp intervals across different library elements to ensure homogenous, full coverage of the template. Either 31 (for *CX3CR1*) or 28 (for both *NRCAM* and *W601*) unique oligo randomized libraries per template (each containing a differently positioned $N_{20}$ tile) were then pooled to create three distinct template-specific libraries (Table S2). These libraries were individually amplified via PCR, purified, verified for size (~147 bp) and measured for concentration, then combined in equimolar amounts to create the 'DNA input library' used in PIONEAR-seq experiments.

For the assembly of nucleosome core particles (NCPs), individual dialysis processes were conducted for each sample (each TF, each NCP type, each round, each replicate) at 4 °C. First, 2 μg of both DNA were mixed in a high-salt buffer (2 M KCl, 10 mM Tris-HCl pH 7.5, 1 mM EDTA, 1 mM DTT) with multiple octamer concentrations to find the optimal DNA:octamer ratio that gave maximal amount of NCP and minimal residual free DNA, and which typically ranged between 1:1.2 and 1:1.8 (Fig. S1B**)**. Then, a peristaltic pump gradually and slowly (~16 hours) lowered the salt concentration, inducing assembly of the NCPs. Next, the samples were dialyzed for 1 hour in low-salt storage buffer (20 mM Tris-HCl pH 7.5, 1 mM EDTA, 1 mM DTT) to stabilize the newly formed NCPs. For each sample, successful NCP assembly was verified using EMSAs (Fig. S1C**)**.

To capture TF-bound NCPs by PIONEAR-Seq assay, each above-described GST-tagged TF was incubated at a concentration of 50 nM with 200 μg of NCP library in 300 μl of PIONEAR–seq binding buffer (10 mM Tris–HCl pH 7.5, 50 mM KCl, 5 mM NaCl, 1 mM $MgCl_2$, 3 μM $ZnSO_4$, 0.5 mM EDTA, 4% glycerol) supplemented with 1 mM DTT, 150 ng poly dI/dC and 40 μl pre-equilibrated Pierce glutathione agarose beads (ThermoFisher, catalog no. 16101) in a low-retention 1.5 ml tube and incubated overnight at 4 °C on a rotatory mixer (~10 second per rotation) as previously described for PIONEAR-seq experiments (*1*). Beads were washed 10 times to remove nonspecifically associated DNA and then used in a PCR reaction to amplify the DNA bound to the TF proteins captured on the beads.

After the initial round of selection (Round 1), each TF-specific PIONEAR-seq DNA library was PCR-amplified and separately assembled onto histone octamers containing the same Histone octamer type of the initial round. The resulting NCPs were evaluated by EMSA to check that the residual free DNA in the samples was also minimal and then incubated in the same conditions of the initial rounds to enrich for the nucleosome DNA sequences with high affinity for the assayed TF. This process was repeated for three rounds by using the DNA amplified in the previous round as the starting material for the next. In parallel, two controls were assayed: (1) a positive control using the libraries as free DNA to check for the ability of the TFs to specifically bind the DNA library; (2) a negative control using GST alone to quantify the enrichment of sequences (see below) in PIONEAR-seq assays.

To prepare samples for Illumina DNA sequencing, the two independent replicates of the Round 3 and 4 libraries of the 7 TFs (as well as the GST control) selected in the 4 NCP conditions (unmodified H3, H3K9me3, H3K27me3, H3K27ac) as well as in the free DNA control were PCR amplified separately using the distinct primer pairs to isolate each template-specific library (*CX3CR1, NRCAM*, W601). The three template-specific libraries obtained from the same sample were considered as 'template replicates' of each other since they underwent the PIONEAR-seq selection process as a pool.

Then, each template-specific library was PCR-amplified as previously described (*1*) to add Illumina adaptors, Unique Molecular Identifiers (UMIs), TruSeq dual-barcodes for multiplexing and P5/P7 promoters for Illumina sequencing (Table S1). The barcoded libraries were quantified (Invitrogen Qubit 4 Fluorometer), verified for the presence of a unique ~300-bp DNA band (Agilent D1000 TapeStation), pooled together to obtain ~$3\times10^6$ sequencing reads for each TF library and ~15×$10^6$ reads for each of the GST and input libraries, and sent for sequencing (HiSeq 4000, 150-bp paired-end) to Azenta (previously Genewiz).

PIONEAR-seq data analysis for k-mer enrichment

Analysis of PIONEAR-seq sequencing reads was carried out essentially as described previously (*1*), as briefly summarized below.

First, for each dataset, the sequencing reads were aligned to the template of origin to find the position of the random tile. The paired-end sequencing reads (forward (FW) and reverse complementary (RC) orientation) were combined into single sequences using PEAR (*6*). To remove potential PCR biases in the amplification process, the UMIs were used to eliminate duplicates (if two reads had the same DNA sequence and the same UMI, they were treated as a single original molecule and the duplicate was discarded). After deduplication, the 6-bp UMI tags were removed, which left reads with an expected length equal to the 147 bp of the template of origin. Reads with length different than 147 bp were dismissed. To extract the DNA sequence and the location of the tiles, the reads were aligned to the associated template using the R function 'pairwiseAlignment' (R package Biostring) with settings: type='overlap', gapOpening=1000, gapExtension=0. Alignments with a score less than 127 were dismissed. By looking at where the DNA sequence differed from the original template (mismatches), the exact position of the $N_{20}$ random tile was identified. Reads were then organized into subsets based on the position of the random tiles (*i.e.*, for the W601 template this resulted in 28 different subsets). As previously reported, a bias that excessively duplicated two 20-mers of the W601-based library, namely GCTTAAACTATGCGCATAGC in the 55–74 bp tile and

ACTATGCGCATAGCTGTCCC in the 61–80 bp tile, was eliminated by removing these elements from the W601-based samples (*1*).

Then, each dataset was normalized to ensure that each tile subset was equally represented ('isotiling'). For that, the median number of reads across all tile subsets in the dataset was calculated and each subset was randomly resampled to contain a number of reads equal to the median, therefore creating a standardized version of the data where each tile position has equal weight. To ensure data reliability, isotiled datasets with fewer than 50K elements were discarded.

Finally, the isotiled datasets were analyzed for *k*-mer enrichment to identify the specific DNA sequence that each TF preferred to bind. To avoid biases coming from the template sequences flanking the random tiles, 2 bp from each end of the 20-bp tiles were removed, which left the 16-bp internal sequence of the tiles for analysis. Each dataset that went through TF PIONEAR-Seq (foreground set) was then compared against the background set represented by its associated initial DNA input library (background set). This comparison was performed by using the workflow of *k*-mer enrichment provided by the SELEX R package (*7, 8*). More precisely, for the most frequent *k*-mers in each dataset, the SELEX R package calculated how many times they appeared in the foreground set (Observed Counts) and how many times they should have appeared if the foreground set were indistinguishable from the background set (Expected Counts). The enrichment of a *k*-mer, which is defined by the ratio of its observed to its expected counts, measures the fold change (FC) of *k*-mer frequency upon TF selection; for the purpose of this analysis, a high ratio indicates a strong binding preference for the *k*-mer by the given TF. To assign the optimal *k*-mer length to each TF (10-mers for OCT4, 9-mers for FOXA1, and 8-mers for CEBPB, GATA1/4, PAX7, RUNX2 and GST mock controls) the values defined in a previous PIONEAR-seq study that used the same TF expression vectors, the same protocol for *in vitro* TF expression and the same DNA libraries were used (*1*). Given the optimal *k*-mer length of each TF, the R package SELEX (*7, 8*) efficiently infers the frequency of *k*-mers of various lengths (typically from 6 to 12 bp) in the background set by using a $5^{th}$-order Markov Model that mitigate the effects of short-range library biases and count sparsity for long *k*-mers. The SELEX package also quantifies the information gain of the library for a given length *k*, which is the sum of the enrichment of all the enriched *k*-mers of length *k* in the dataset weighted by each *k*-mer's frequency and is formally similar to the Kullback-Leibler divergence between the *k*-mer frequency distribution in the selected library versus the input library (*7, 9*). The most enriched *k*-mer (Fig. 1B, left panel) and the information gain (Fig. 1B, right panel) of each dataset are provided in Table S4.

<u>Analysis of 8-mer positional preference in PIONEAR-seq data</u>

Analysis of 8-mer positional preference in PIONEAR-seq data was carried out essentially as described previously (*1*). Briefly, the absolute positional preference profiles of specific 8-mers in each sample of interest were calculated by first counting their matches within the isotiled 16-bp random regions at template position. By convention, the 8-mer coordinates referred to the position of their first base relative to the 147-bp template sequence oriented from 5' to 3' end. Depending on the template's primers, the 8-mer positional preference profiles exhibited slightly different ranges: from bp 20 to bp 118 in *CX3CR1*-based, from bp 28 to bp 117 in *NRCAM*-based and from bp 24 to bp 110 in W601-based libraries. The absolute positional preference profiles were divided by the total number of 8-mer counted matches in the dataset to create normalized positional preference profiles, which can then be compared with each other.

AlphaFold3 models of TF:nucleosome binding

AlphaFold3 (AF3) from the AF3 server (alphafoldserver.com) (*10*) was used to model the structures analyzed for the template modeling score (Fig. 3G; Fig. 4E) and for the solvent-accessibility surface area (Fig. 4I). Representative structures were used in (Fig. 3C, D, H; Fig. 4D, I; Fig. S7, Fig. S8, Fig. S10). Details of the amino acids and DNA sequences used as AF3 input can be found in Table S5 and the full reports generated by AF3 for each model can be found in the Harvard Dataverse repository https://doi.org/10.7910/DVN/RMT39Y. Each report includes the 5 predicted structures that AF3 modeled from the diffusion network obtained with the same initial seed by adding random noise during the final coordinate generation. In all the AF3 models for nucleosomes, two copies of the four unmodified full-length *Xenopus laevis* histones (H2.A, H2.B, H3, H4) were used. Since both the PAX7-specific HD.2 motif and the FOXA1-specific DIV0 motif are palindromic and represent DNA-mediated dimerization for the associated TFs, 2 copies of the TF's DBD were included in the input of the AF3 models. For PAX7, solely the HD part of the DBD was used to model binding to the HD.2 motif. Both for PAX7 and FOXA1 binding, AF3 models of their binding to 14-bp free DNA molecules containing the TF-specific motifs for PAX7 (TATAATCGATTATA, Fig. S8**)** and for FOXA1 (AGTAAATATTTACT, Fig. S10B) were generated to create reference structures of DNA-mediated TF dimerization. For both PAX7 and FOXA1, AF3 models of their interactions with nucleosomes were generated by tiling the internal region of the 147-bp W601 DNA sequence with the above-mentioned motifs, so that the PAX7's HD.2 8-mer AATCGATT appeared at each internal position between bp 31 and 110 (used for the analysis in Fig. 3G) and the FOXA1's DIV0 8-mer AAATATTT appeared at each internal position between bp 26 and 105 (used for the analysis in Fig. 4E). To evaluate the background recruitment of PAX7 and FOXA1 to W601 nucleosomes, AF3 models for their binding to a nucleosome containing an unmodified 147-bp W601 sequence were also generated and used to create the dashed lines in the analysis in Fig. 3G and Fig. 4E, respectively. In the structural models presented in Fig. 3H and Fig. 4D, PyMol was used to add the H3 tail (a.a. residues 24-43) of an NCP solved by cryo-EM (PDB: 8G8G) (*11*) onto the indicated AF3 models by aligning the globular part (command 'super') of the respective H3 histones.

<u>DNA-mediated TF dimerization on NCP</u>

To evaluate which sites of the nucleosomal DNA allow the HD.2 and the DIV0 motifs to mediate the dimerization of, respectively, PAX7 and FOXA1, each AF3 model of TF-bound nucleosomes (a few examples in Fig. S7 for PAX7 and in Fig. S10A for FOXA1) was compared to the associated AF3 model of TF-bound free DNA (Fig. S8 for PAX7 and Fig. S10B for FOXA1), which exhibited the correct DNA-mediated TF dimerization. The template modeling (TM) score provided by the recently developed tool US-align (*12*) was used to quantify the alignment of the TF-bound free DN to the TF-bound nucleosome. Since the TF-bound free DNA should align to just the portion of the nucleosome at the TF binding site, US-align was used in multimer mode (-mm 1 -ter 1), to find the best chain mapping between the two complexes. For each AF3 TF-bound nucleosome model, the 5 AF3 predicted structures were separately assessed by US-align and the best TM score was assigned to that model, thus representing how well the model's motif position within the nucleosome was able to correctly accommodate the TF binding (Fig. 3G for PAX7 and Fig. 4E for FOXA1). As negative controls for the motif insertion on the DNA-mediated dimerization of the TFs, the models for W601 with no HD.2 and no DIV0 were also calculate, thus representing how well the TFs could dimerize onto a W601 nucleosome that did not contain the cognate motif (dashed lines in Fig. 3G for PAX7 and Fig. 4E for FOXA1).

Solvent-accessibility surface area analysis

AlphaFold3 (AF3) from the AF3 server (alphafoldserver.com)(*10*) was used to model a nucleosome containing the 147-bp W601 DNA sequence and the two copies of the four unmodified full-length *Xenopus laevis* histones used also to generate the AF3 models for TF:nucleosome binding in this study (Table S5). The full report for the AF3 model includes the 5 predicted structures that AF3 generated by adding random noise to the diffusion network and can be found in the Harvard Dataverse repository https://doi.org/10.7910/DVN/RMT39Y.

For each structure: (1) the Shrake-Rupley Solvent-accessibility surface area (SASA) (*13*) was computed for each residue of the whole nucleosome by using the Bio.PDB.SASA (biopython package bio.PDB); (2) then an 8-mer window was slid along the two antiparallel DNA strands in the internal single-gyre region around the dyad (bp 74) from bp 60 to bp 88, and the 8 residue bases of the forward strand (*i.e.*, bp 60-67) were matched to their reverse complement bases (*i.e.*, bp 81-88); (3) and finally, the SASA of each window was calculated as the sum of the per-residue SASA over those 16 residues (8 per strand). For the whole AF3 model, the SASA of each 8-mer window was then quantified by the median SASA value of the 5 predicted structures (Fig. 4H).

Motif analysis of TF-bound nucleosome sequences in K562 cells

The hg19 genome will be the reference genome used in the following analysis. 3 bam files of K562 MNase-seq experiments were downloaded from ENCODE (*14*) under the accession codes ENCFF000VMJ, ENCFF000VNO and ENCFF000VNQ and indexed using SAMtools (v1.19) (*15*). Nucleosome footprints were called using DANPOS3 with the parameter setting: -m 0 --frsz 140 -q 5 -jw 40 -jd 100 -z 20 -a 10 -f 0. The nucleosome footprints of length 140 bp (‘K562-nuclesomes') were filtered for further analysis (*16*).

Optimal irreproducible discovery rate (IDR)-thresholded ChIP-seq peaks for CEBPB (accession code: ENCFF813LOW) and OCT4 (accession code: ENCFF990CFV) in K562 cells were downloaded from ENCODE (*14*) in narrowPeak bed format (‘TF-bound-peaks’). While the downloaded OCT4 peaks were aligned to hg19, the downloaded CEBPB peaks were aligned to GRCh38 and their genomic coordinates were converted into hg19 coordinates with liftover (*17*). For each TF, the Intersect function from BEDTools (v2.31.0) (*18*) was used to select the nucleosomes that overlap at least 1 bp of a TF-bound peaks with parameter setting: -a K562-nuclesomes -b TF-bound-peaks -u > TF-bound-nucleosomes. This step created the ‘CEBPB-bound-nucleosomes’ file and the ‘OCT4-bound-nucleosomes’ file, which will be analyzed in parallel in the following. The DNA sequences of TF-bound-nucleosomes were scanned for the presence of the TF motif (at least one instance) with MEME/FIMO (v5.5.9) with option ‘--thresh’ set equal to 1e-4. For CEBPB-bound-nucleosomes the ChIP-Seq-derived motif MA0466.1 and for OCT4-bound nucleosomes the ChIP-Seq-derived motif MA1115.2 from the JASPAR 2024 CORE vertebrate motif database (*19*) were used. The presence of the TF motif was used to label TF-bound nucleosomes that are directly recruiting the chipped TF.

H3K27ac ChIP-seq peaks in K562 cells were downloaded from ENCODE (*14*) in bed narrowPeak format (aligned to hg19) under the accession code ENCFF044JNJ (‘H3K27ac-peaks’). To distinguish between H3K27ac+ and H3K27ac- nucleosomes among TF-bound-nucleosomes, the Intersect function from BEDTools (v2.31.0) (*18*) was used to with parameters setting: -a TF-bound-nucleosomes -b H3K27ac-peaks -c . This step used the overlap with H3K27ac ChIP-seq peaks to create both a set of 'TF-bound H3K27ac+ nucleosomes' and a set of 'TF-bound H3K27ac- nucleosomes'. The amount of TF-bound H3K27ac+ and H3K27ac- nucleosomes that contained the TF binding motif were then retrieved and compared with a Fisher exact test (Fig. 4J).

**Figure S1. Generation of PIONEAR-seq NCP with modified H3**

(**A**) FPLC size-exclusion chromatogram of the *in vitro* assembly of distinct histone octamers, color coded according to the histone 3 types. The fractions collected for subsequent NCP assembly are highlighted by the top segments. The chromatograms are representative of multiple independent replicates. (**B**) EMSA gels for the *in vitro* NCPs reconstituted at the indicated ratio of histone octamer versus DNA input library. The optimal DNA:octamer ratio is indicated in bold. This experiment was repeated independently twice with similar results. (**C**) Nucleosome assembly in subsequent rounds of PIONEAR-seq selection. The images are representative of gels obtained for the any rounds of selection and using different TFs and different types of histone 3.

**Figure S2. Motif enrichment analysis of PIONEAR-seq data for replicates of the 5 TFs that do not show difference in presence of H3K27me3**

(**A**) Data from an independent replicate of the experiments in Fig. 1C. The enrichment of each *k*-mer is measured as the fold change (FC) of its frequency upon TF selection, which is the ratio between the normalized counts observed in the PIONEAR-seq sample (*i.e.*, Round 4) and the normalized counts observed in the input library (Round 0). The normalized counts observed in the input library represents the count frequency expected in the sample if the TF is not selecting the *k*-mer and is estimated by using a $5^{th}$-order Markov model (see Methods). (**B**, **C**) Data from the indicated template replicates of the experiments in Fig. 1C, and representative of two independent replicates. The enrichment of each *k*-mer is quantified as the fold change (FC) of its frequency after selection for binding by a TF, as defined in panel A and in Fig. 1B, C.

**Figure S3. Motif enrichment analysis of PIONEAR-seq data obtained with the indicated H3 types for 5 TFs that do not show difference in presence of H3K27me3**

(**A**, **B**) Same analysis as in (Fig. 1C), but for PIONEAR-seq experiments performed with histone octamer containing either H3K9me3 (A) or H3K27ac (B). The plots are representative of two independent replicates on the pool of the 3 template libraries, namely *CX3CR1* (depicted here), *NRCAM* and W601. The enrichment of each *k*-mer is quantified as the fold change (FC) of its frequency in the TF-selected library versus the Input library, as defined in Fig. 1B and Fig. S2A.

**Figure S4. Motif enrichment analysis of FOXA1 PIONEAR-seq replicates**

(**A**, **B**) Data from an independent replicate of the experiments in (Fig. 2B, C). (**C**–**F**) Data from the indicated template replicates (C-D for *NRCAM*, E-F for W601) of the experiments in (Fig. 2B, C) and representative of two independent replicates. The enrichment of each *k*-mer is quantified as the fold change (FC) of its frequency in the TF-selected library versus the Input library, as defined in Fig. 1B and Fig. S2A.

**Figure S5. Motif enrichment analysis of PAX7 PIONEAR-seq replicates**

(**A**, **B**) Data from an independent replicate of the experiments in (Fig. 2E, F). (**C**–**F**) Data from the indicated template replicates (C, D for *NRCAM*, E, F for W601) of the experiments in Fig. 2E, F, and representative of two independent replicates. The enrichment of each *k*-mer is quantified as the fold change (FC) of its frequency in the TF-selected library versus the Input library, as defined in Fig. 1B and Fig. S2A.

**Figure S6. Analysis of the positional preference of PAX7 binding in PIONEAR-seq data**

(**A**) Data from an independent replicate of the experiments in Fig. 3A. (**B**, **C**) Data from the indicated template replicates of the experiments of Fig. 3A, both representative of two independent replicates. The heatmap for the HD.2 motif in panel C is displayed also in the left panel of Fig. 3E.

**Fig. S7. Predicted structures for PAX7 binding on W601-based NCPs by AlphaFold3**

AlphaFold3-modeled (AF3) structures of 2 PAX7 HD DBDs and W601 nucleosomes with an HD.2 8-mer at the indicated positions. Structures with HD.2 starting at bp 34 and 45 same as in Fig. 3C, D, respectively. The AF3 model structure with HD.2 starting at bp 66 was used for the structural alignment presented in Fig. 3H. PAX7 HDs are in cyan, histones in green, and DNA backbones in gray except for HD.2 8-mers, which are in red.

**Figure S8. AlphaFold3 predicted structures of PAX7 binding to HD.2 free DNA**

AlphaFold3 (AF3) structural model for the interaction between 2 PAX7 HD DBDs and an HD.2 8-mer inserted in the middle of a 14-bp free DNA molecule (TATAATCGATTATA). Colors as in Fig. S7.

**Figure S9. Analysis of the positional preference of FOXA1 binding in PIONEAR-seq data**

(**A**) Data from an independent replicate of the experiments in Fig. 4A. (**B**) Data from template replicates of the experiments in Fig. 4B, each depicting two independent replicates (**C**) Heatmaps of the density profile of 8-mers DEG2 (both in forward *(left)* and reverse complementary *(middle)* orientations) and DIV0 (right), for two independent replicates. The heatmaps of the DIV0 motif on the right appear also in Fig. 4B.

**Figure S10. AlphaFold3 predicted structures of FOXA1 binding to W601-based nucleosomes**

(**A**) AlphaFold3 models of the interaction between 2 FOXA1 DBDs and nucleosomes assembled with 147-bp W601 sequences that contain the DIV0 8-mers at the indicated positions. Model for DIV0 starting at bp 71 was also used to generate Fig. 4D. FOXA1 DBDs are in cyan, histones in green, DNA backbones in gray except for the DIV0 8-mers, which in red. (**B**) AlphaFold3 model for the interaction between 2 FOXA1 DBDs and the DIV0 8-mer inserted in the middle of a 14-bp free DNA molecule (AGTAAATATTTACT). Colors as in (A). Three images of the same structural model are presented.

**Fig. S11. Analysis of the positional preference of CEBPB and OCT4 in PIONEAR-seq data**

Data from an independent replicate of the experiments in Fig. 4F.
